# scRNA-seq analysis of hemocytes of penaeid shrimp under virus infection

**DOI:** 10.1101/2023.01.04.521844

**Authors:** Keiichiro Koiwai, Hidehiro Kondo, Ikuo Hirono

## Abstract

The classification of cells in non-model organisms has lagged behind that of model organisms using established cluster of differentiation marker sets. To reduce fish diseases, research is needed to better understand immune-related cells, or hemocytes, in non-model organisms like shrimp and other marine invertebrates. In this study, we used Drop-seq to examine how virus infection affected the populations of hemocytes in kuruma shrimp, *Penaeus japonicus*, which had been artificially infected with a virus. The findings demonstrated that virus infection reduced particular cell populations in circulating hemolymph and inhibited the expression of antimicrobial peptides. We also identified the gene sets that are likely to be responsible for this reduction. Additionally, we identified functionally unknown genes as novel antimicrobial peptides, and we supported this assumption by the fact that these genes were expressed in the population of hemocytes that expressed other antimicrobial peptides. In addition, we aimed to improve the operability of the experiment by conducting Drop-seq with fixed cells as a template and discussed the impact of methanol fixation on Drop-seq data in comparison to previous results obtained without fixation. These results not only deepen our understanding of the immune system of crustaceans but also demonstrate that single-cell analysis can accelerate research on non-model organisms.

## Introduction

Less is known about invertebrate immune responses than about mammalian immune responses. This is due to the fact that invertebrates, or non-model organisms, have not yet had their marker genes, or the cluster of differentiation markers (CD markers), thoroughly reported as they are in mammals. Invertebrates are currently responsible for the vast majority of aquaculture production worldwide, and it is anticipated that their significance will continue to grow (Naylor et al., 2021). In order to prevent diseases and develop resources for human food, it is crucial to understand how invertebrate immune systems work. In many nations around the world, aquaculture is used to produce penaeid shrimp, such as *Penaeus chinensis, P. japonicus, P. monodon*, and *P. vannamei*. However, diseases always affect a shrimp aquaculture farm, so effective disease-controlling techniques are required to minimize the harm (Flegel, 2019). Penaeid shrimp are invertebrates, meaning they lack antibody-mediated adaptive immunity and are thought to only have innate immunity to pathogens. Therefore, it is essential to research how hemocytes, which are circulating in body fluids, respond to pathogens during an infection with a disease because they control innate immunity.

Understanding cell functions and physiological roles starts with classification. For cell classification, population variation, and functional analysis in humans and other mammals, CD markers for cell surface proteins have been developed. It is challenging to develop markers that are equivalent to mammal CD markers in non-model organisms with a small number of researchers because these CD markers are the result of the development of diverse monoclonal antibodies by many researchers over a long period of time. The development of short-read next-generation sequencers has enabled techniques to be developed in the last ten years or so to thoroughly analyze the mRNA from a single-cell (scRNA-seq), regardless of the targeting of every eukaryote. With the aid of microfluidic devices, techniques like Drop-seq (Macosko et al., 2015), inDrop (Klein et al., 2015), HyDrop (De Rop et al., 2022), and Chromium (10x Genomics) have successfully increased the number of analyzed cells to be in the thousands or more as opposed to the hundreds as in the case of cell sorters. In previous years, the mRNA expression of whole tissues or populations of cells has been examined; however, these scRNA-seq techniques now make it simple to examine differences in gene expression between individual cells. These techniques are also useful for classifying cell populations in tissues without the use of markers because they enable the analysis of large numbers of single cells at once. Previously, authors used Drop-seq to classify hemocytes of *P. japonicus*, which had primarily been classified by morphology, and they proposed new cell populations and markers for these populations (Koiwai et al., 2021). The analysis is currently being conducted on aquaculture species like penaeid shrimp (Cui et al., 2022; Li et al., 2022; Yang et al., 2022; Zhu et al., 2021), crayfish (Söderhäll et al., 2022), and other shellfish (Meng et al., 2022; Meng & Wang, 2022; Sun et al., 2021). Future efforts to control diseases in aquaculture species will benefit from knowing this information. The majority of the results, however, are only based on the analysis of healthy individuals because scRNA-seq is still prohibitively expensive and unsuitable for handling large numbers of samples.

White spot syndrome disease (WSSD), whose causative agent is white spot syndrome virus (WSSV), is one of the main diseases that cause issues in crustacean aquaculture (Flegel, 2012). After 30 years since the disease’s initial discovery in the 1990s, there is still no fundamental strategy for controlling it, and it continues to do significant harm year after year. It is important to understand whether WSSV infection suppresses shrimp immune function due to changes in the cell population of hemocytes or whether it results in changes in specific gene expression that make it impossible to control WSSV infection. This knowledge can then be applied to develop novel anti-WSSD control strategies, such as techniques to prevent alterations in the cell population or boost the expression of a particular gene. In order to study whether virus infection alters the hemocyte population or how gene expression changes, scRNA-seq of hemocytes was performed on healthy and WSSV-infected *P. japonicus*. Through this research, we found that virus infection reduced particular cell populations in circulating hemolymph and inhibited the expression of antimicrobial peptides. We also identified the gene sets that are likely to be responsible for this reduction, and functionally unknown genes as novel antimicrobial peptides. Additionally, marker genes on hemocytes that could be used in penaeid shrimp with or without viral infection were also found. There were difficulties in terms of technology and timing when preparing cells for scRNA-seq right away after infection testing. Methanol was used to fix the hemocytes used in this study, and fixed hemocytes were kept that way until scRNA-seq was done. The impact of methanol fixation on shrimp scRNA-seq analysis was also discussed by contrasting the results obtained with methanol-fixed hemocytes with those obtained with unfixed hemocytes previously performed.

## Results

### Decreased hemocyte numbers after viral infection

Injections of approximately 10,000 copies of the WSSV into shrimp of the size used in this study led to 100% mortality within 4 days, according to preliminary infection tests. The virus concentration in the gills of moribund shrimps was more than 10^5-6^ copies/ng of total DNA extracted from the gills. Therefore, to estimate the function of hemocytes under the most severe conditions of WSSV infection, we analyzed hemocytes of shrimps by Drop-seq with a WSSV copy number greater than 10^5^ copies/ng of DNA from gills after artificial WSSV infection. Information on the sex, non-infected or infected by WSSV, and WSSV copy number of shrimps subjected to Drop-seq is shown in Table 1. The library was successfully prepared from a total of eight shrimp, two males and two females each of WSSV uninfected and infected individuals. After sequencing these eight libraries, the archived Drop-seq results of hemocytes from three *P. japonicus* (Koiwai et al., 2021) were also integrated, and the data from a total of 11 shrimps were subjected to further bioinformatic analysis.

**Table 1.**
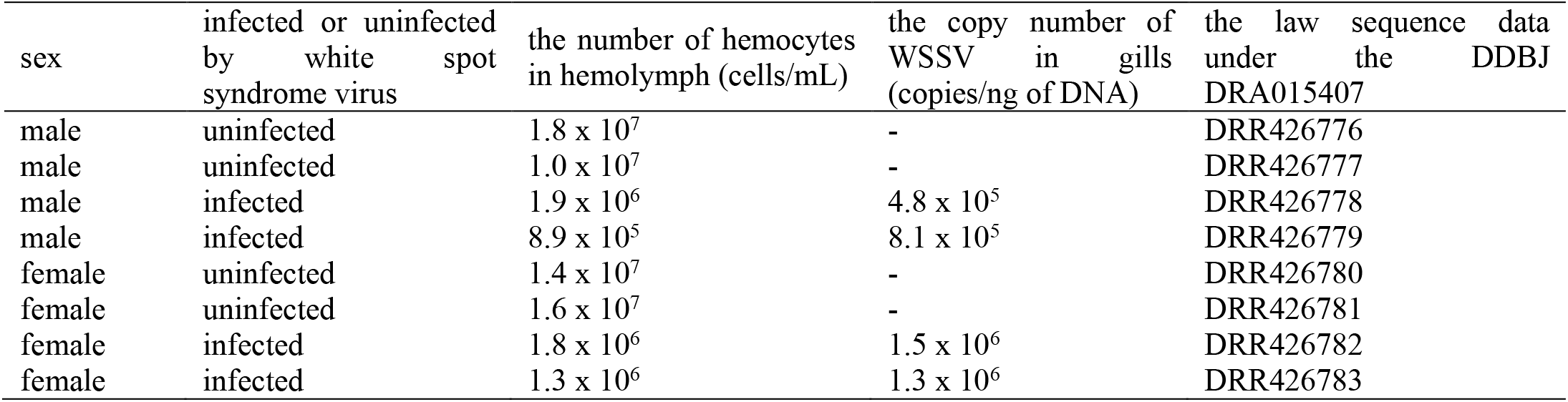
The information of shrimp used for Drop-seq in this experiment

### Impact of methanol fixation on hemocytes single-cell analysis

Single-cell gene expression data from a total of 6,788 hemocytes from 11 shrimp were dimensionally compressed into a uniform manifold approximation and projection (UMAP) as shown in Figure 1A. The number of clusters for a set resolution parameter of 0.5 was eight, tentatively named Hem1-8, respectively. The shape of the UMAP was similar to the previous results (Koiwai et al., 2021), suggesting that the Drop-seq libraries constructed in this experiment were successfully prepared and sequenced. The median number of mRNA counts (UMI number) per cell was 852, the median number of genes detected was 443, and the median percentage of mitochondria-derived genes expressed was 1.77% (Figure 1 B-D). The protocol using methanol-fixed hemocytes showed a decrease in the number of detected UMIs per cell from 1,012 to 805 and genes per cell from 515 to 498, and an increase in the percentage of mitochondria-derived genes expressed from 1.77% to 1.90%, compared to using fresh hemocytes (Figure 1-figure supplement 1A-C). This is similar to existing reports that scRNA-seq with methanol fixation has a poor detection rate when compared to living cells (Alles et al., 2017; X. Wang et al., 2021). On the other hand, there was no bias in the UMAP figure, indicating that Seurat’s SCTransform function was successfully integrated (Figure 1-figure supplement 2).

**Figure 1.**
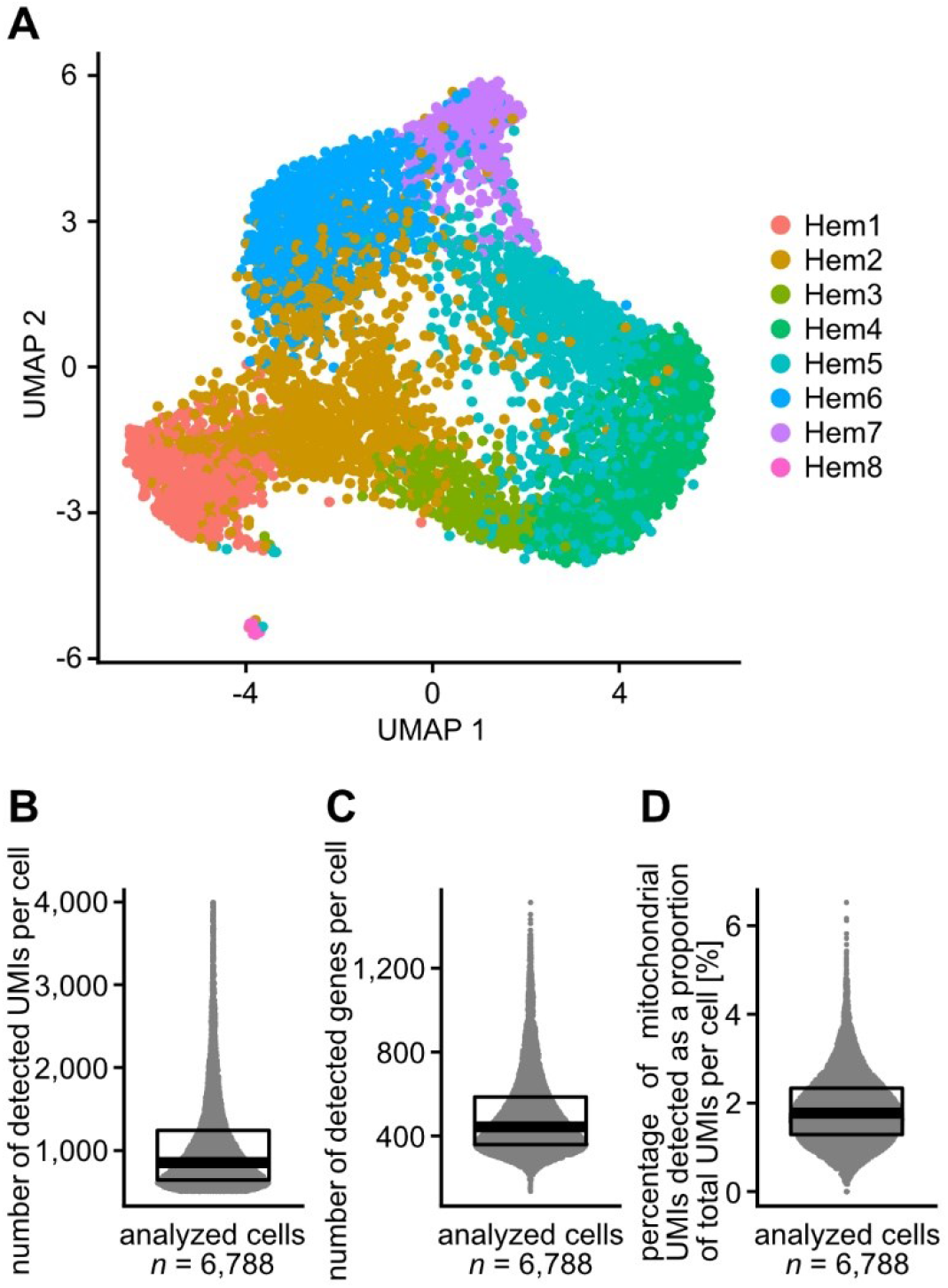
Hemocyte single-cell analysis from 11 shrimp. The uniform manifold approximation and projection (UMAP) of 6,788 hemocytes of *Penaeus japonicus* (A). The number of detected mRNA counts (UMI counts) per cell (B), the number of genes per cell (C), and the expressed percentage of mitochondria-derived genes (D). The median values of UMIs per cell, genes per cell, and percentage of mitochondria-derived genes were 852, 443, and 1.77%, respectively.

**Figure 1-figure supplement 1.**
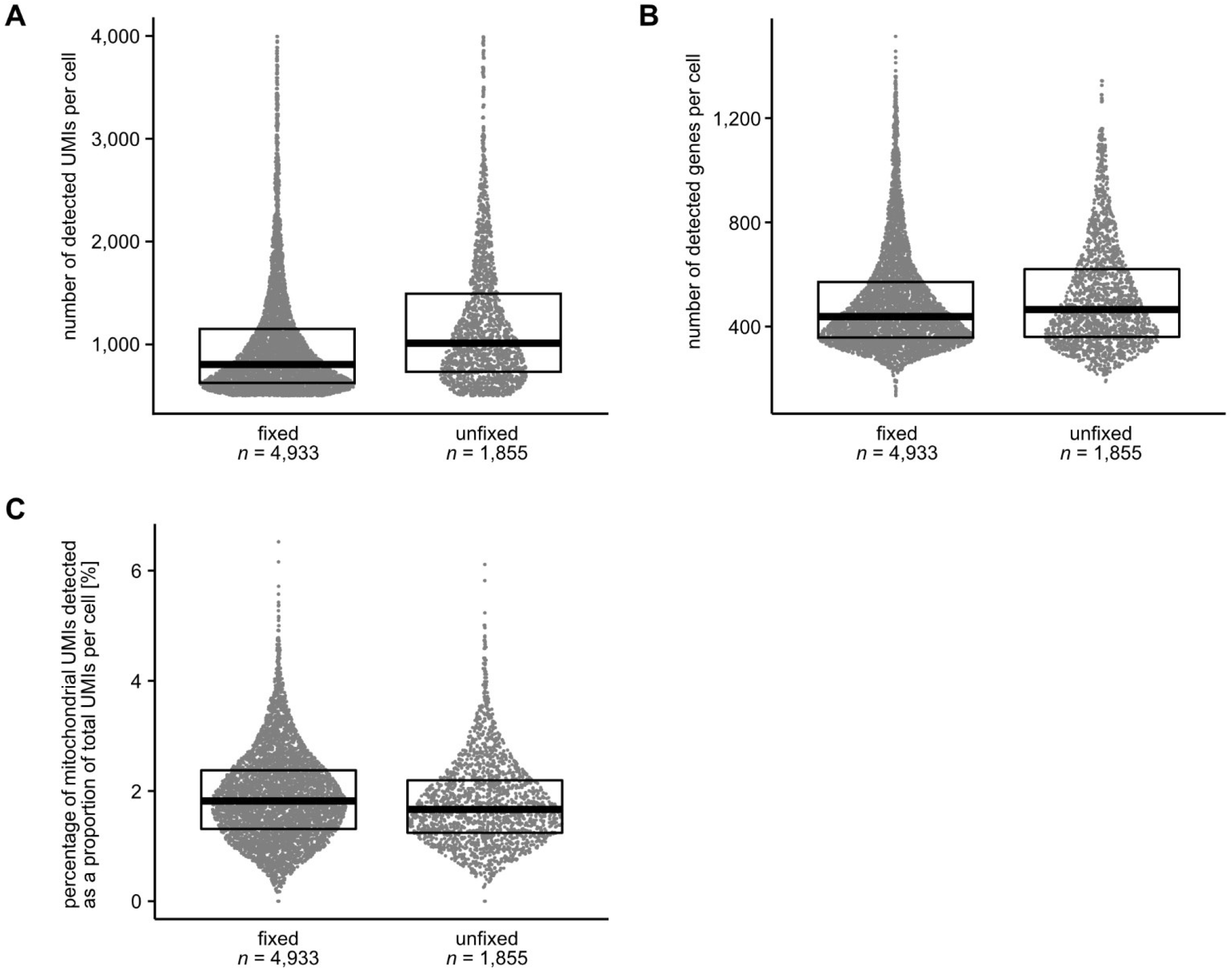
Comparison with the fixed procedure of detection sensitivity on hemocytes. The median values of the number of detected mRNA counts (UMI counts) per cell were 805 for fixed and 1,012 for unfixed (A). The median number of detected gene counts per cell was 498 for fixed and 515 for unfixed (B). The median percentages of expressed mitochondria-derived genes were 1.90% for fixed and 1.77% for unfixed (C).

**Figure 1-figure supplement 2.**
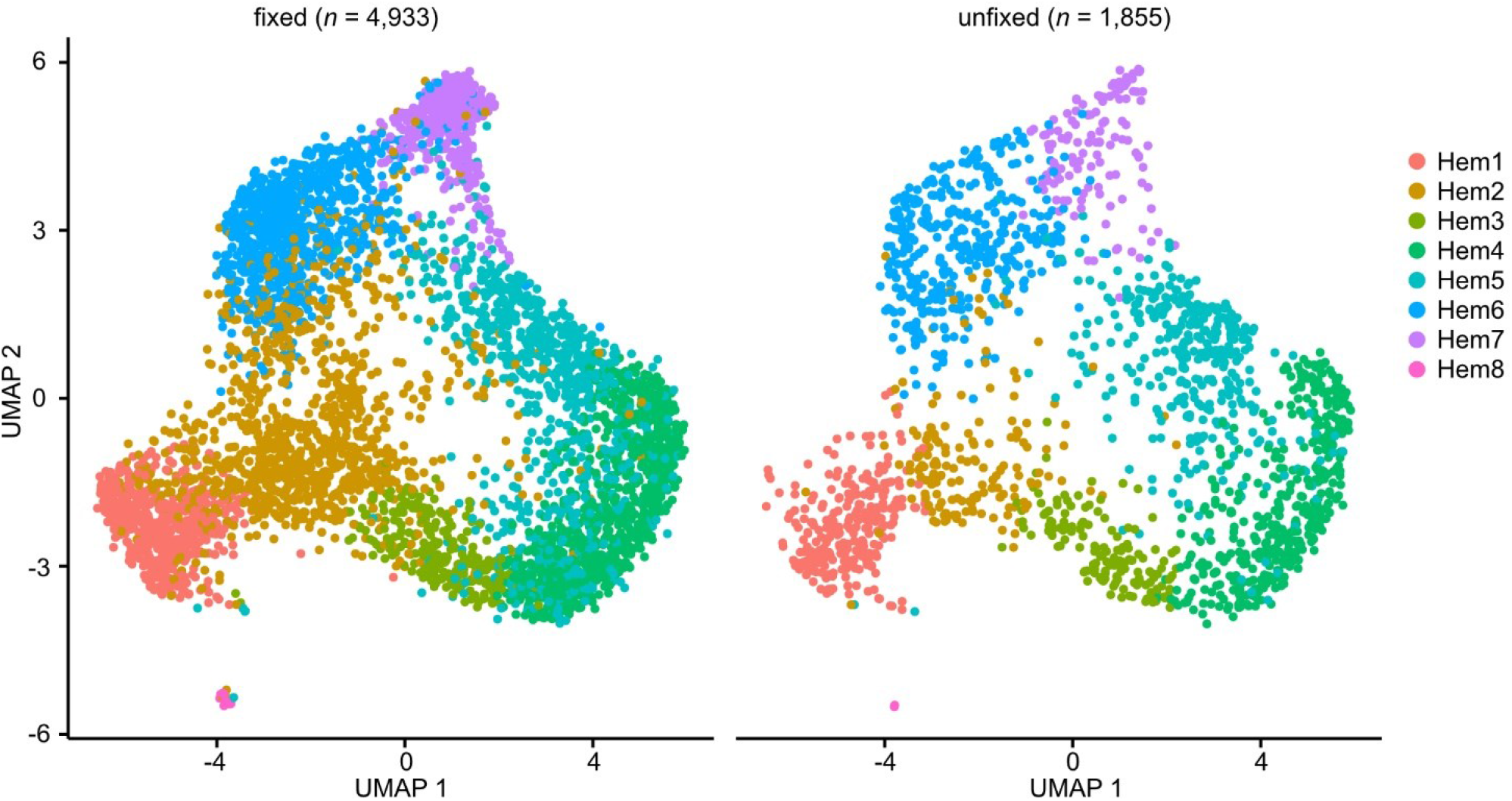
Comparison with the fixed procedure of uniform manifold approximation and projection (UMAP). UMAP from fixed and unfixed hemocytes of *P. japonicus*.

### Hemocyte cluster variation caused by WSSV infection

In shrimp infected with WSSV, the percentage of clusters Hem2 and Hem8 increased while Hem4 decreased (Figure 2AB). WSSV infection reduces the number of hemocytes in body fluids (Cui et al., 2020; Elbahnaswy et al., 2017; van de Braak et al., 2002; Wongprasert et al., 2003). Therefore, we calculated the actual number of each clustered hemocyte in the body fluid, not the percentage of each cluster. Infection with WSSV reduced the number of hemocytes in all clusters (Hem1–7) except Hem8 (Figure 2C). These results indicate that WSSV infection not only decreases any type of hemocyte but also specifically decreases the Hem4 cluster. The percentage of expressed genes from WSSV was detected in each cluster, and cluster Hem7 had a higher proportion of WSSV-derived mRNA than the other clusters (Figure 2-figure supplement 1). Additionally, the cluster Hem8 was excluded from downstream analysis because it is not sufficient in number for further analysis.

**Figure 2.**
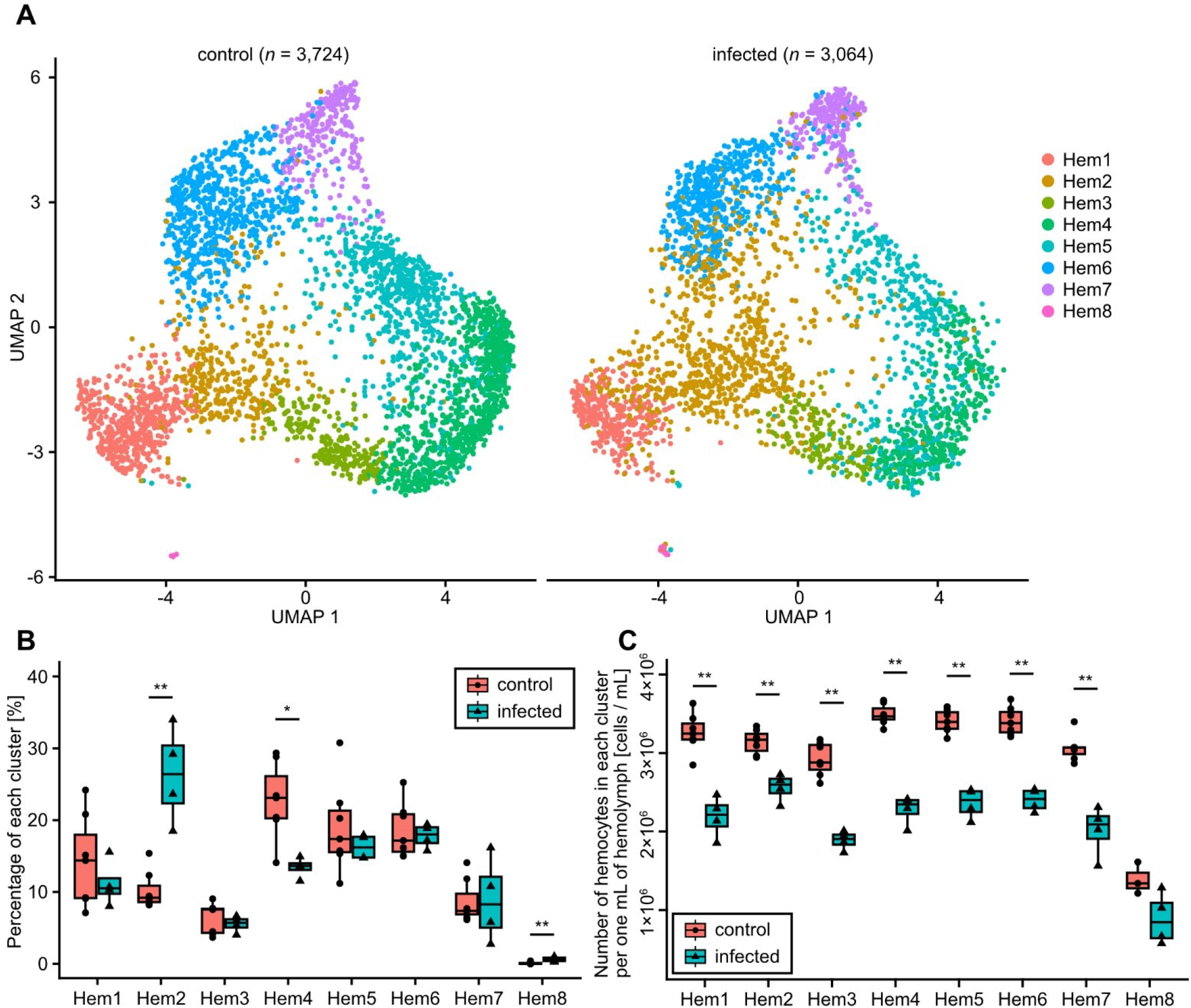
Changes in hemocyte clusters due to WSSV infection. The uniform manifold approximation and projection (UMAP) from the control and infected groups (A). The percentage (B) and the actual number (C) of hemocytes in each cluster. The significance between the control and infected groups was calculated by the Wilcoxon signed-rank test. The *p* values shown in the figures are represented by \**p < 0*.*05 and* \*\**p* < 0.01.

**Figure 2-figure supplement 1.**
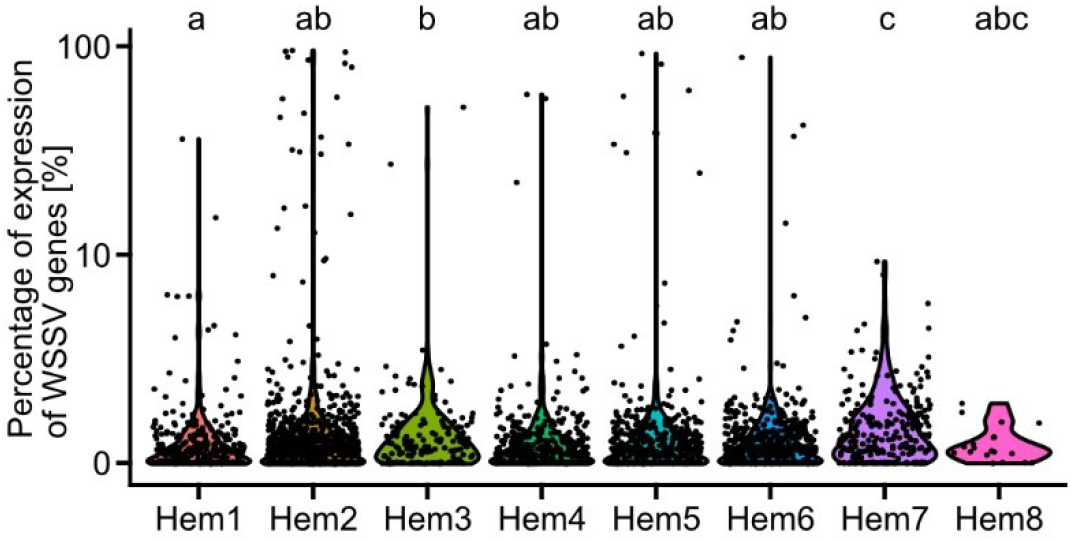
The percentage of expressed genes from WSSV in each cluster. One-way ANOVA was conducted, with letters above each cluster to show which groups are statistically different from one another by *p*<0.05.

### Differentially expressed genes caused by WSSV infection in total hemocytes or each cluster

How WSSV infection affected gene expression in hemocytes in circulating hemolymph was investigated. First, the genes that varied more than 2-fold in more than 50% of all hemocytes were extracted (Figure 3). As a result, six genes whose expression level is increased and 11 genes whose expression level is decreased by WSSV infection were detected. Also, genes that changed 2-fold or more in more than 50% of all hemocytes were extracted specifically for each cluster (Figure 3-figure supplement 1-7).

**Figure 3.**
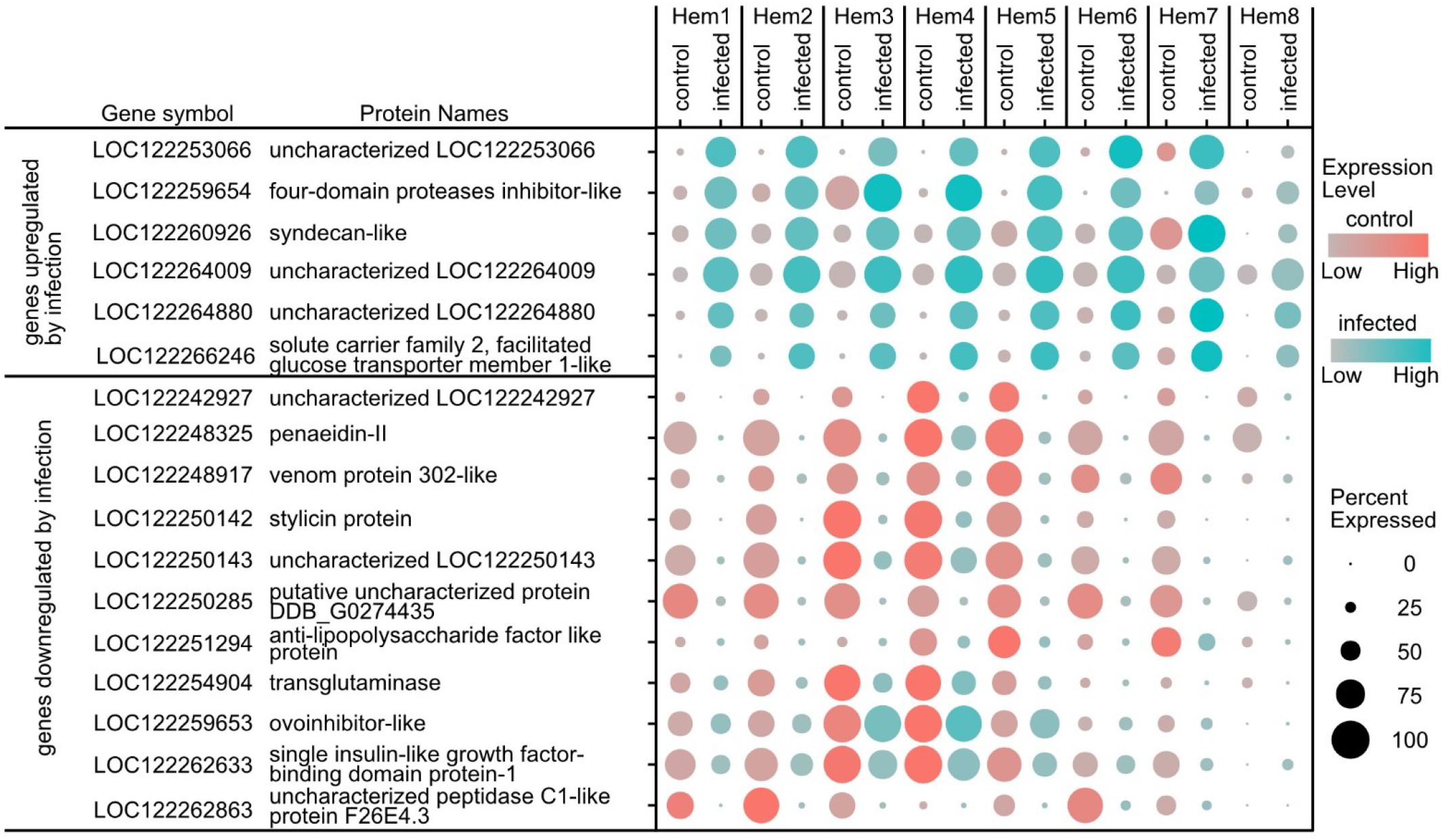
Dot plot profiling of differentially expressed genes caused by WSSV infection in total hemocytes. The color gradient of dots represents the expression level, while the size represents the percentage of cells expressing any genes per cluster.

**Figure 3-figure supplement 1.**
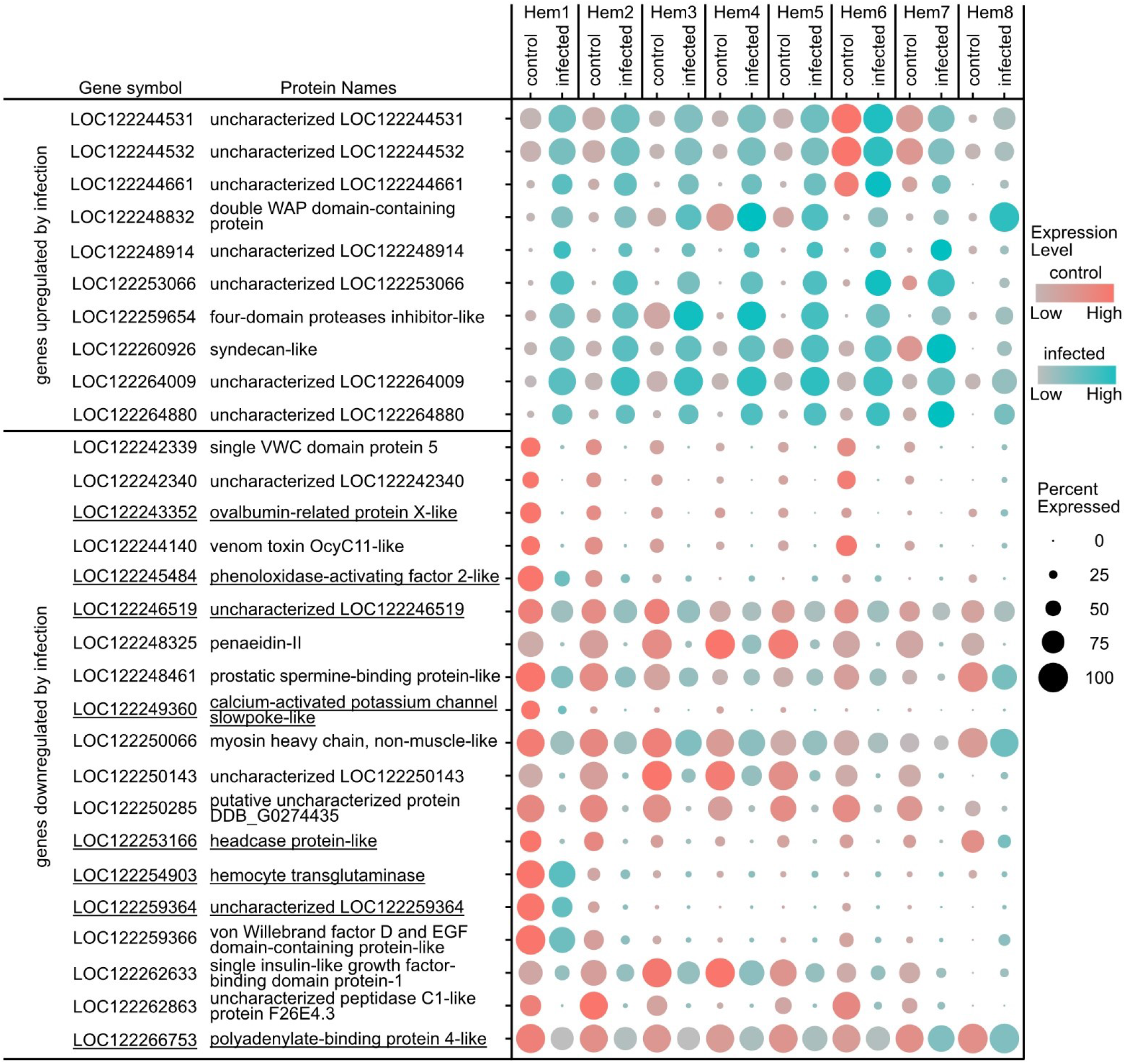
Dot plot profiling of differentially expressed genes caused by WSSV infection in cluster Hem1. The color gradient of dots represents the expression level, while the size represents the percentage of cells expressing any genes per cluster. Underlines indicate genes that were specifically affected in this cluster.

**Figure 3-figure supplement 2.**
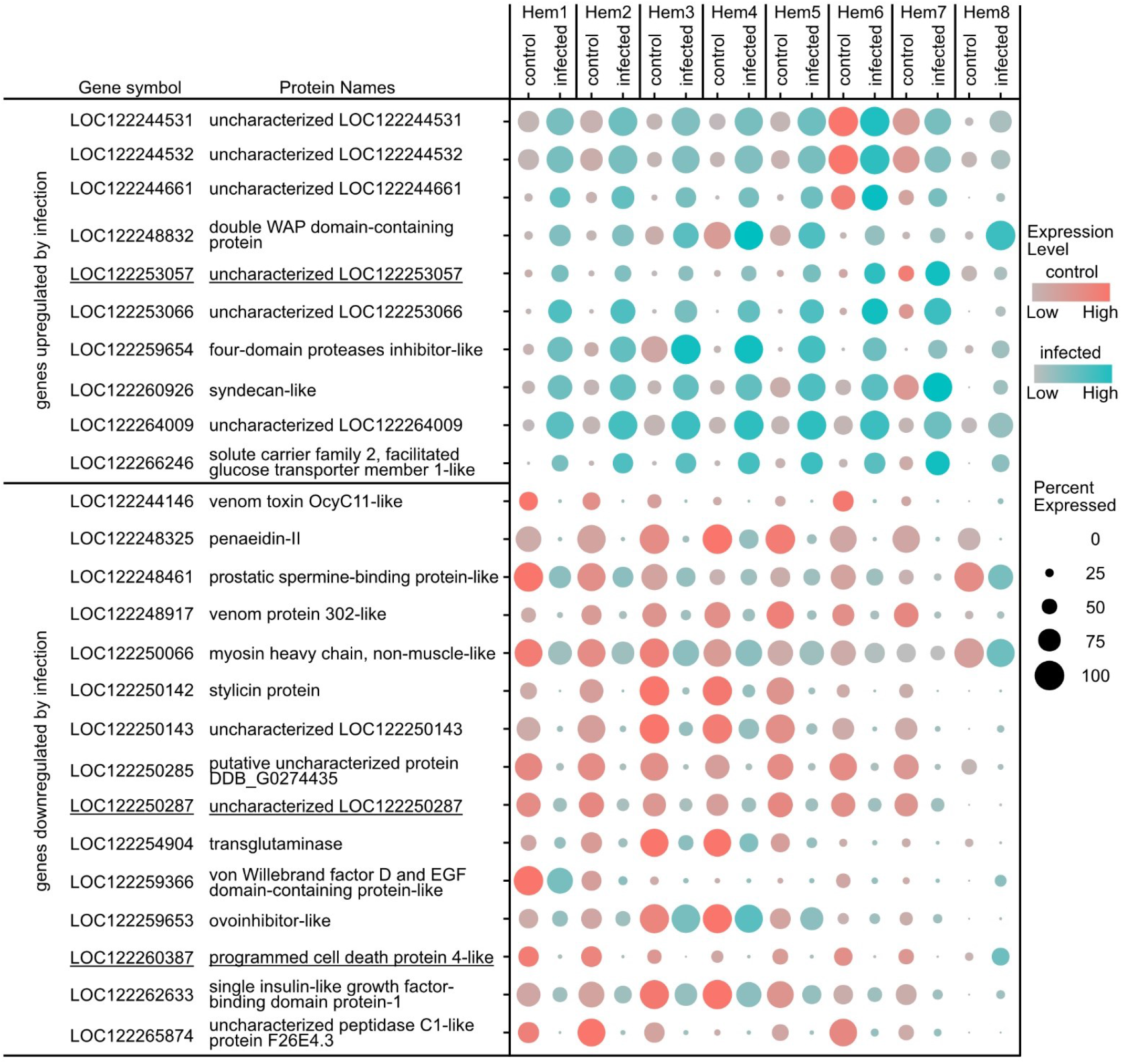
Dot plot profiling of differentially expressed genes caused by WSSV infection in cluster Hem2. The color gradient of dots represents the expression level, while the size represents the percentage of cells expressing any genes per cluster. Underlines indicate genes that were specifically affected in this cluster.

**Figure 3-figure supplement 3.**
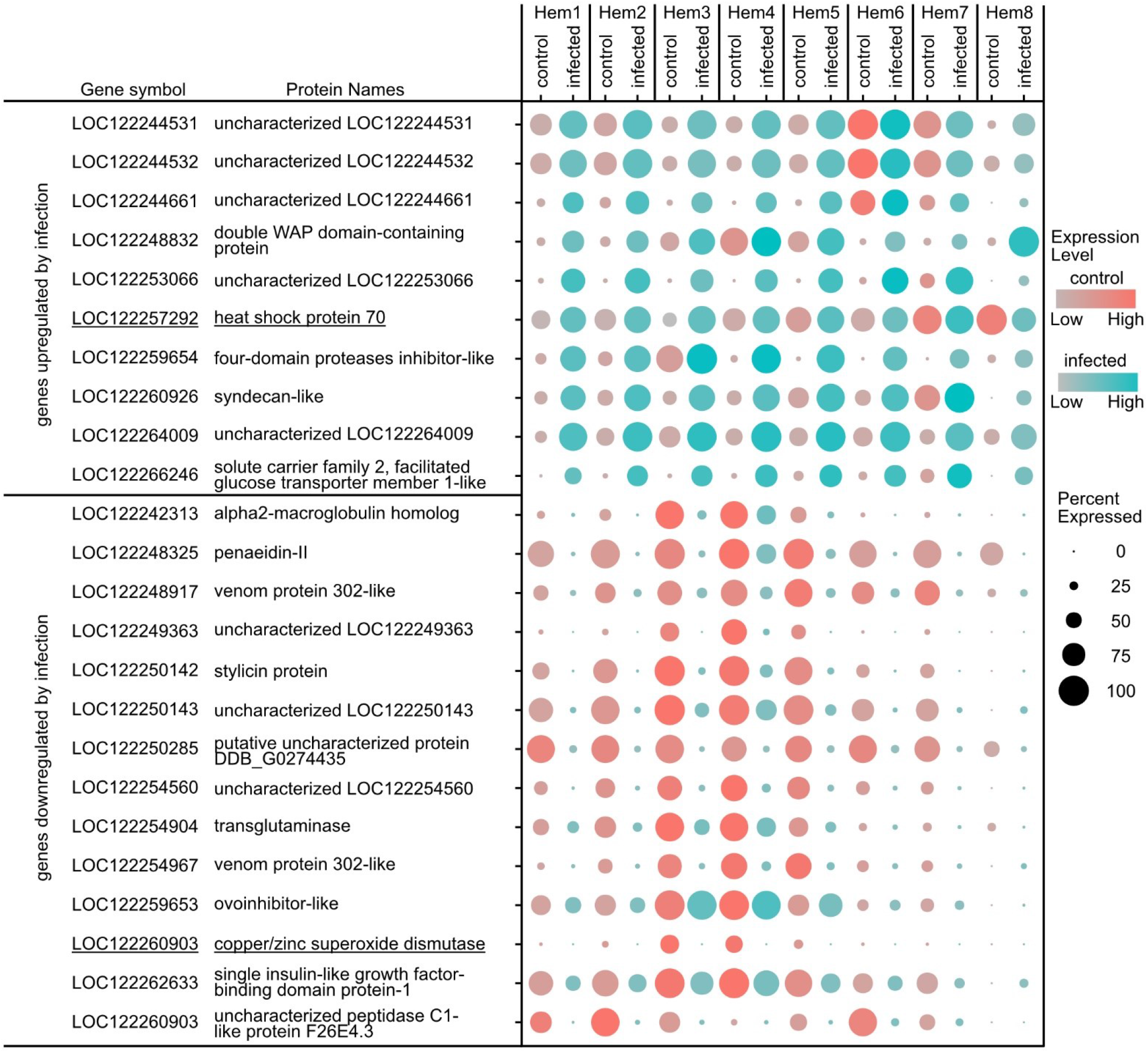
Dot plot profiling of differentially expressed genes caused by WSSV infection in cluster Hem3. The color gradient of dots represents the expression level, while the size represents the percentage of cells expressing any genes per cluster. Underlines indicate genes that were specifically affected in this cluster.

**Figure 3-figure supplement 4.**
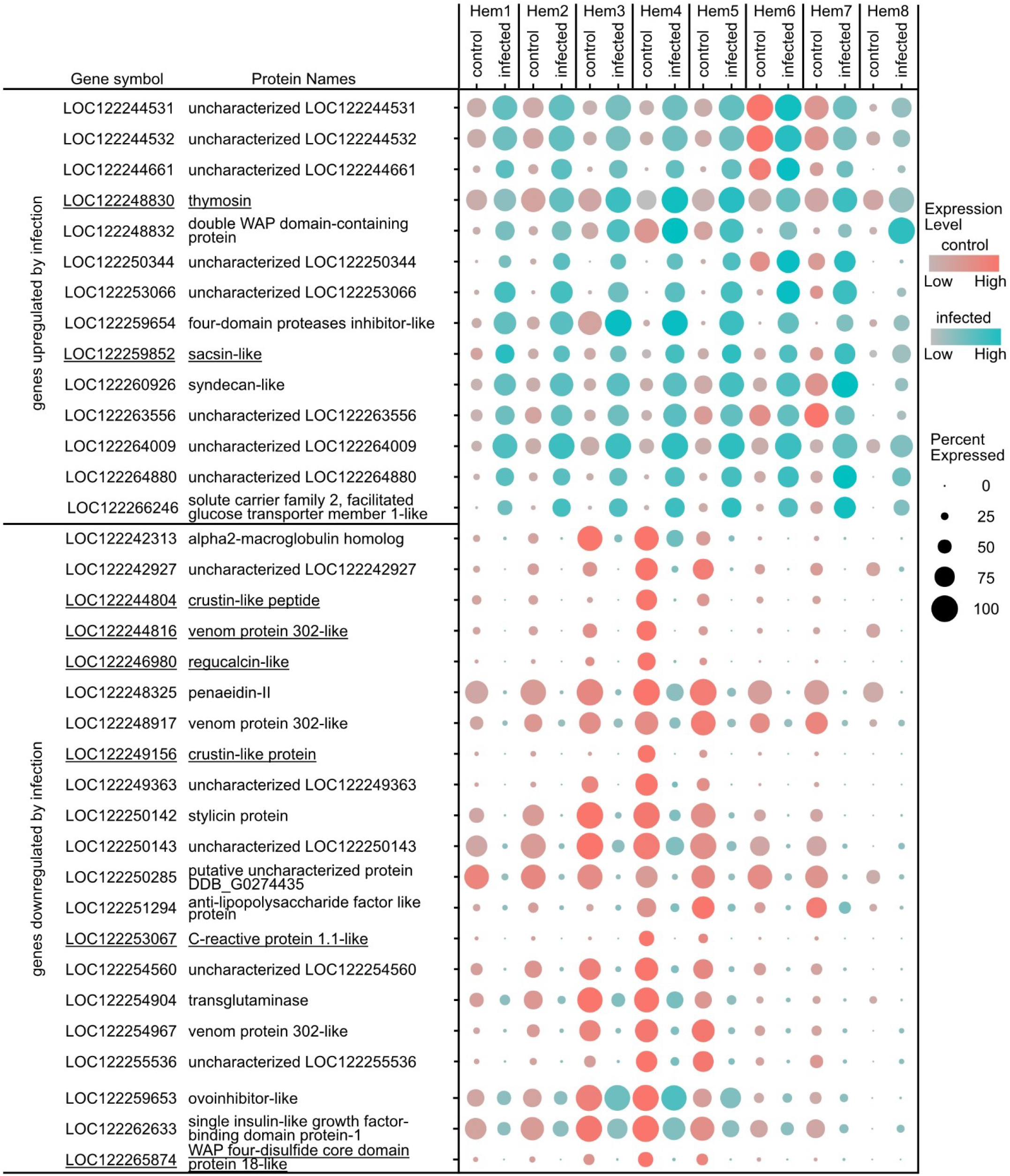
Dot plot profiling of differentially expressed genes caused by WSSV infection in cluster Hem4. The color gradient of dots represents the expression level, while the size represents the percentage of cells expressing any genes per cluster. Underlines indicate genes that were specifically affected in this cluster.

**Figure 3-figure supplement 5.**
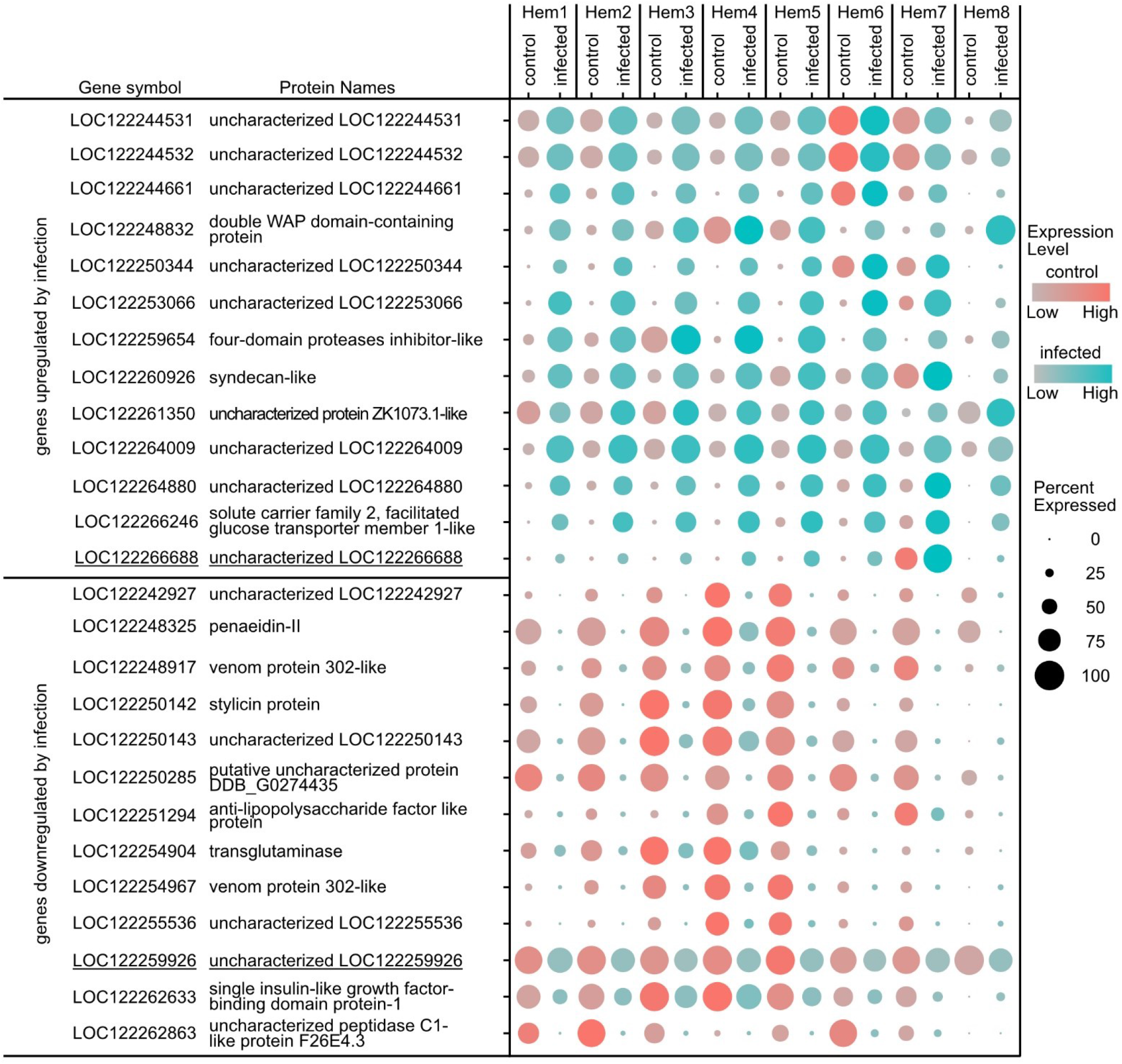
Dot plot profiling of differentially expressed genes caused by WSSV infection in cluster Hem5. The color gradient of dots represents the expression level, while the size represents the percentage of cells expressing any genes per cluster. Underlines indicate genes that were specifically affected in this cluster.

**Figure 3-figure supplement 6.**
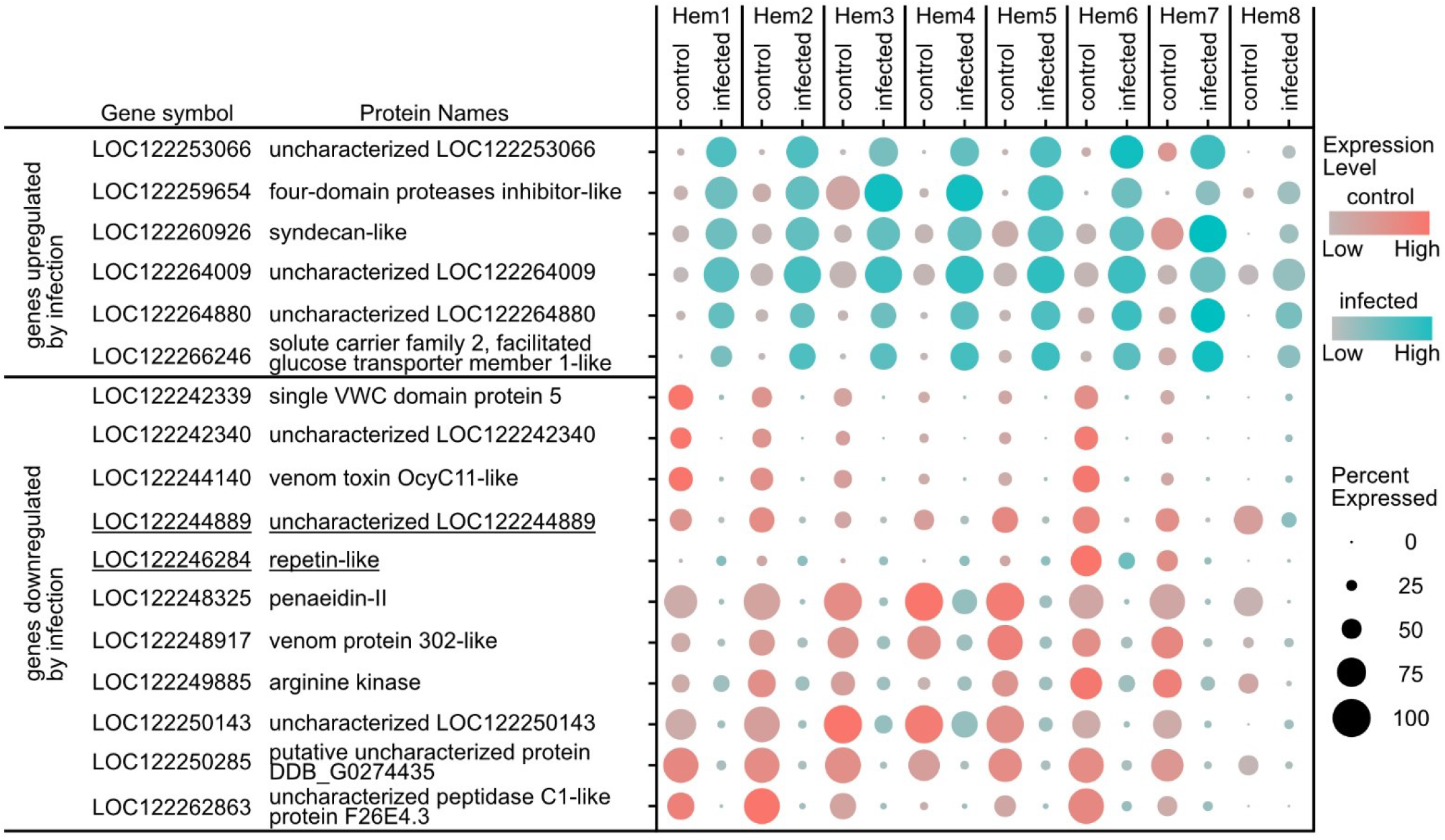
Dot plot profiling of differentially expressed genes caused by WSSV infection in cluster Hem6. The color gradient of dots represents the expression level, while the size represents the percentage of cells expressing any genes per cluster. Underlines indicate genes that were specifically affected in this cluster.

**Figure 3-figure supplement 7.**
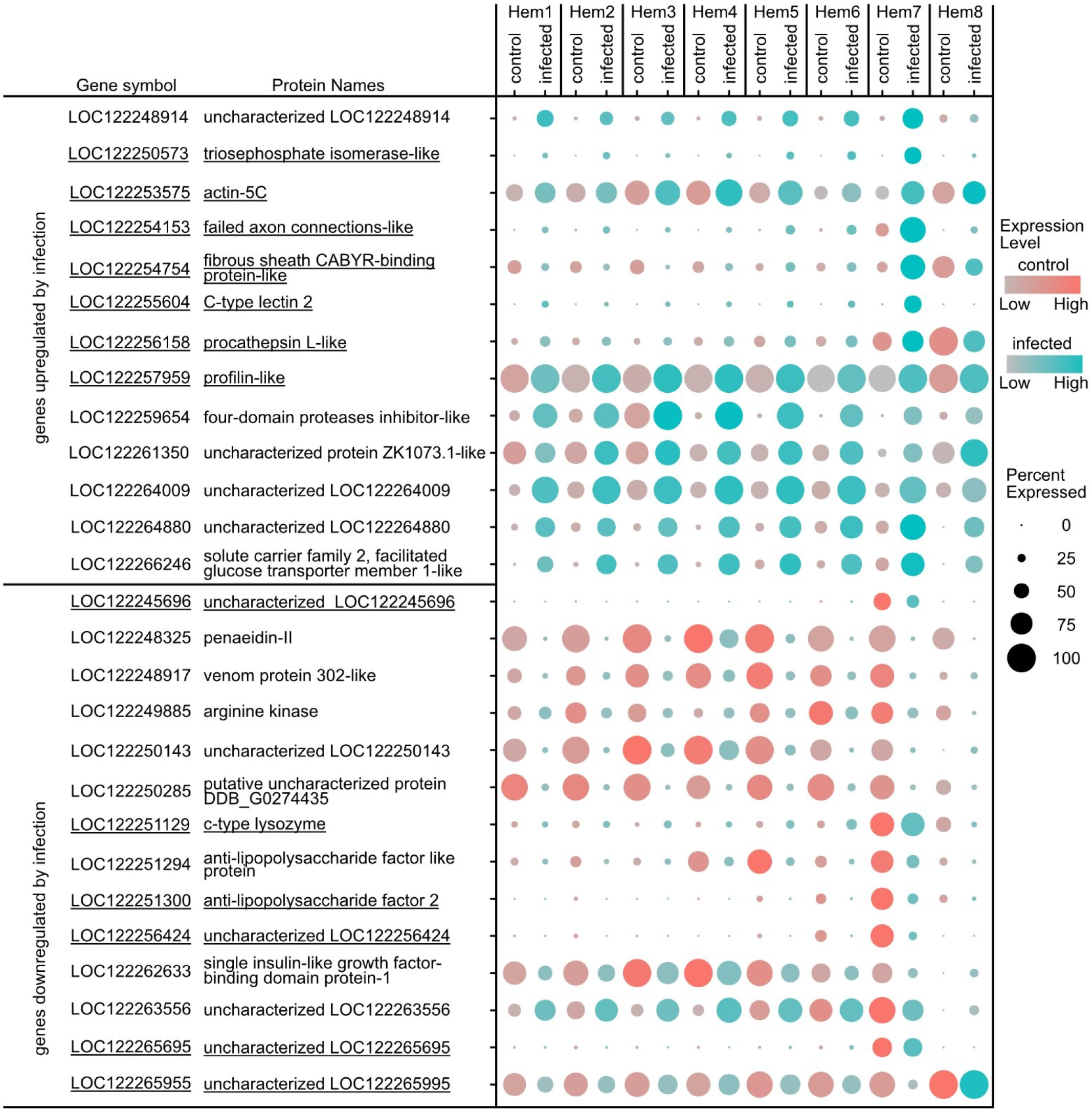
Dot plot profiling of differentially expressed genes caused by WSSV infection in cluster Hem7. The color gradient of dots represents the expression level, while the size represents the percentage of cells expressing any genes per cluster. Underlines indicate genes that were specifically affected in this cluster.

The *venom protein 302-like* genes, LOC122244816, LOC122248917, and LOC122254967, are among those in cluster Hem4 that change during viral infection (Figure 3-figure supplement 4). These genes were predicted to have a single insulin-like growth factor binding protein (IGFBP) domain at the N-terminal end. Through a blast search, crustacean hematopoietic factor (CHF) from a shrimp belonging to the same genus, *P. vannamei*, was hit. LOC122262633 *single insulin-like growth factor binding protein-1* was another gene with a single IGFBP domain that is strongly expressed in cluster Hem4, suggesting proteins with this domain play an important role during viral infection.

### Marker genes that characterize clusters even under virus infection

To establish stable markers to distinguish specific hemocyte clusters, genes whose expression does not change with virus infection are appropriate. Therefore, we predicted marker genes for each cluster with or without WSSV infection as conserved markers. Four genes were predicted as candidate markers for Hem1, five for Hem4, three for Hem6, six for Hem7, and four for Hem8, respectively (Figure 4).

**Figure 4.**
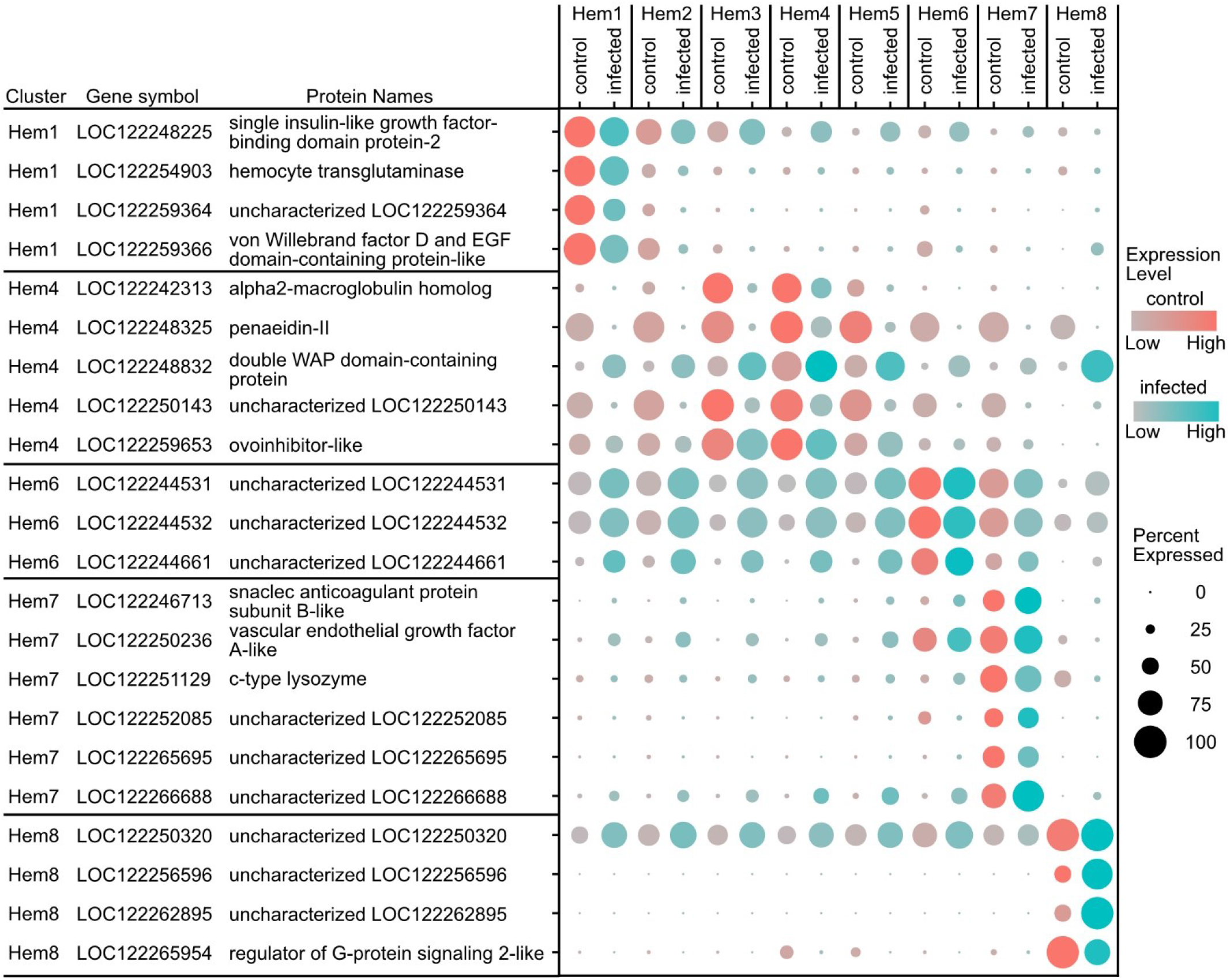
Dot plot profiling of conserved marker genes for each cluster of hemocytes with or without WSSV infection. The color gradient of dots represents the expression level, while the size represents the percentage of cells expressing any genes per cluster.

**Figure 4-figure supplement 1.**
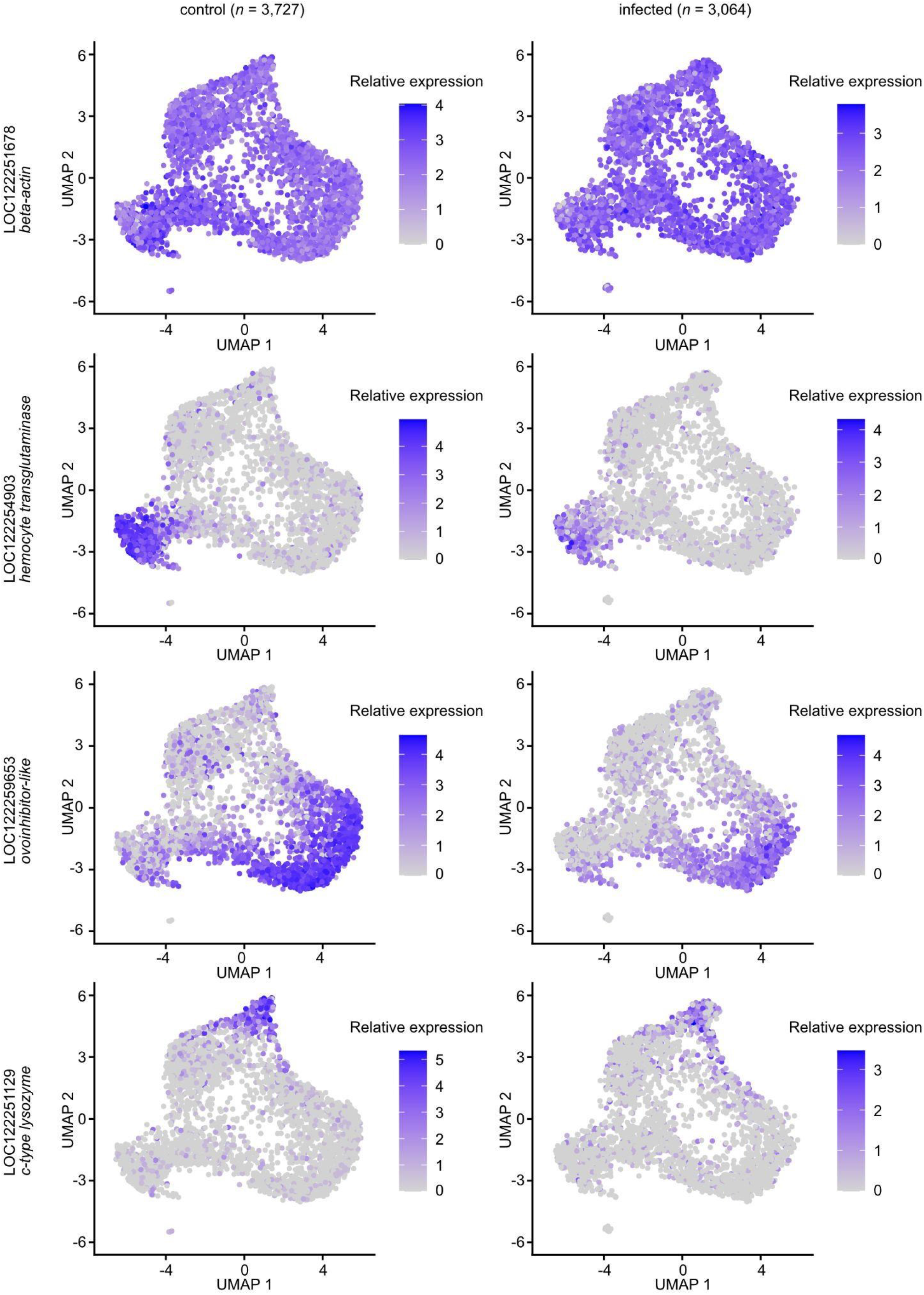
Expression pattern of three marker genes in comparison with the fixed procedure of uniform manifold approximation and projection (UMAP). The color gradient of the bars represents the expression level. A housekeeping gene, LOC122251678 *beta-actin*, is uniformly expressed in all clusters, whereas marker genes are expressed in a cluster-specific manner. The expression levels of each marker gene were not affected by viral infection.

**Figure 4-figure supplement 2.**
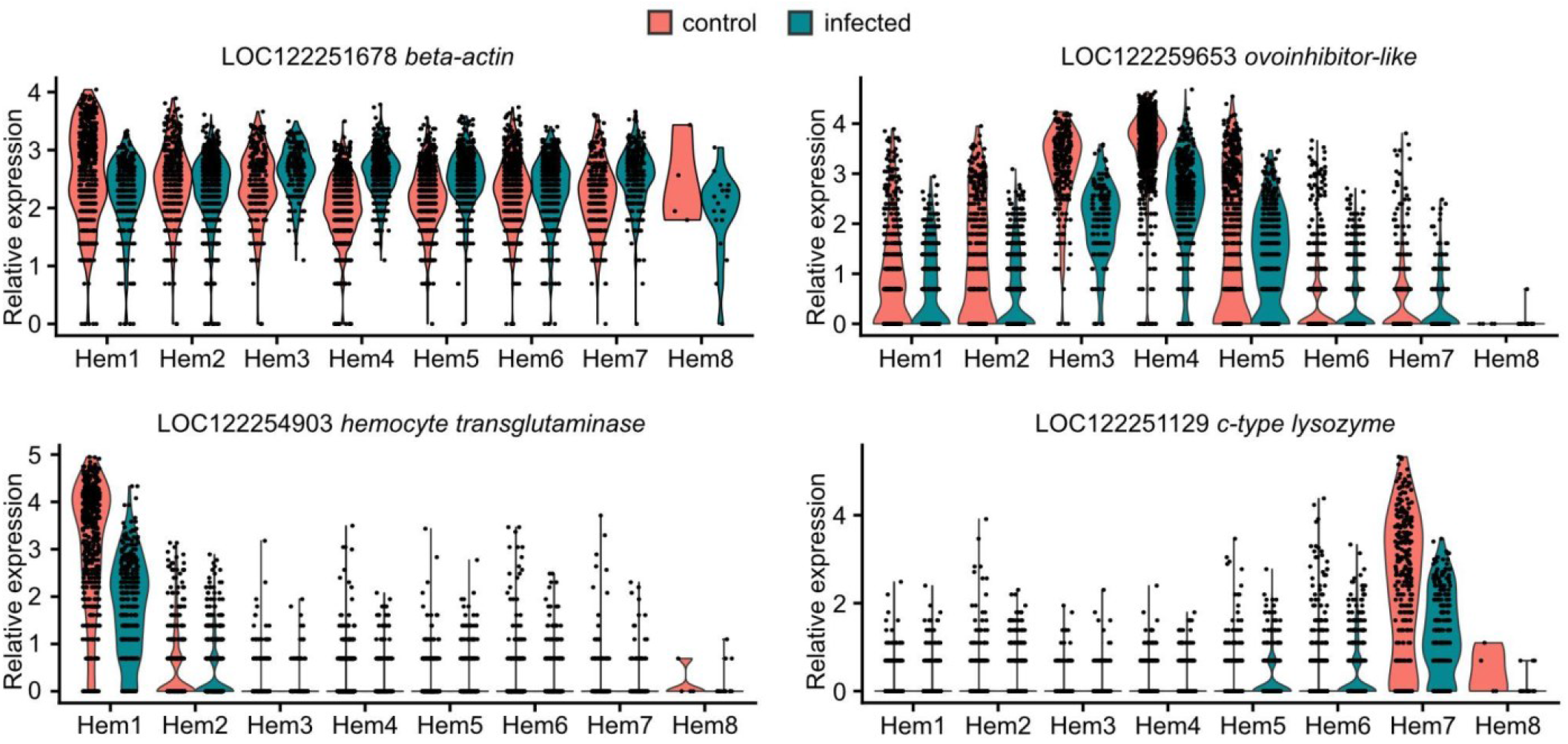
Visualization of expression distributions of three marker genes in each cluster. A housekeeping gene, LOC122251678 *beta-actin*, is uniformly expressed in all clusters, whereas marker genes are expressed in a cluster-specific manner. The expression levels of each marker gene were not affected by viral infection.

The predicted marker genes for cluster Hem1, Hem4, and Hem7, whose clusters are discussed as the origins and two endpoints of differentiation, were detected by single molecule fluorescence *in situ* hybridization (smFISH). Three genes were selected for smFISH observation, LOC122254903: Hem1, LOC122259653: Hem4, and LOC122251129: Hem7 (Figure 4-figure supplement 1-2). Every gene was expressed only in specific hemocytes (Figure 5AB), and furthermore, LOC122259653 *ovoinhibitor-like* and LOC122251129 *c-type lysozyme* were never transcribed in the same cell (*n* = 306 individual cells). The mRNA was clearly transcribed in each hemocyte without any reduction due to viral infection. The positive percentage of LOC122254903 was 10.4%: control; 9.3%: infected, LOC122259653 was 42.0%: control; 64.6%: infected, and LOC122251129 was 11.1%: control; 6.3%: infected.

**Figure 5.**
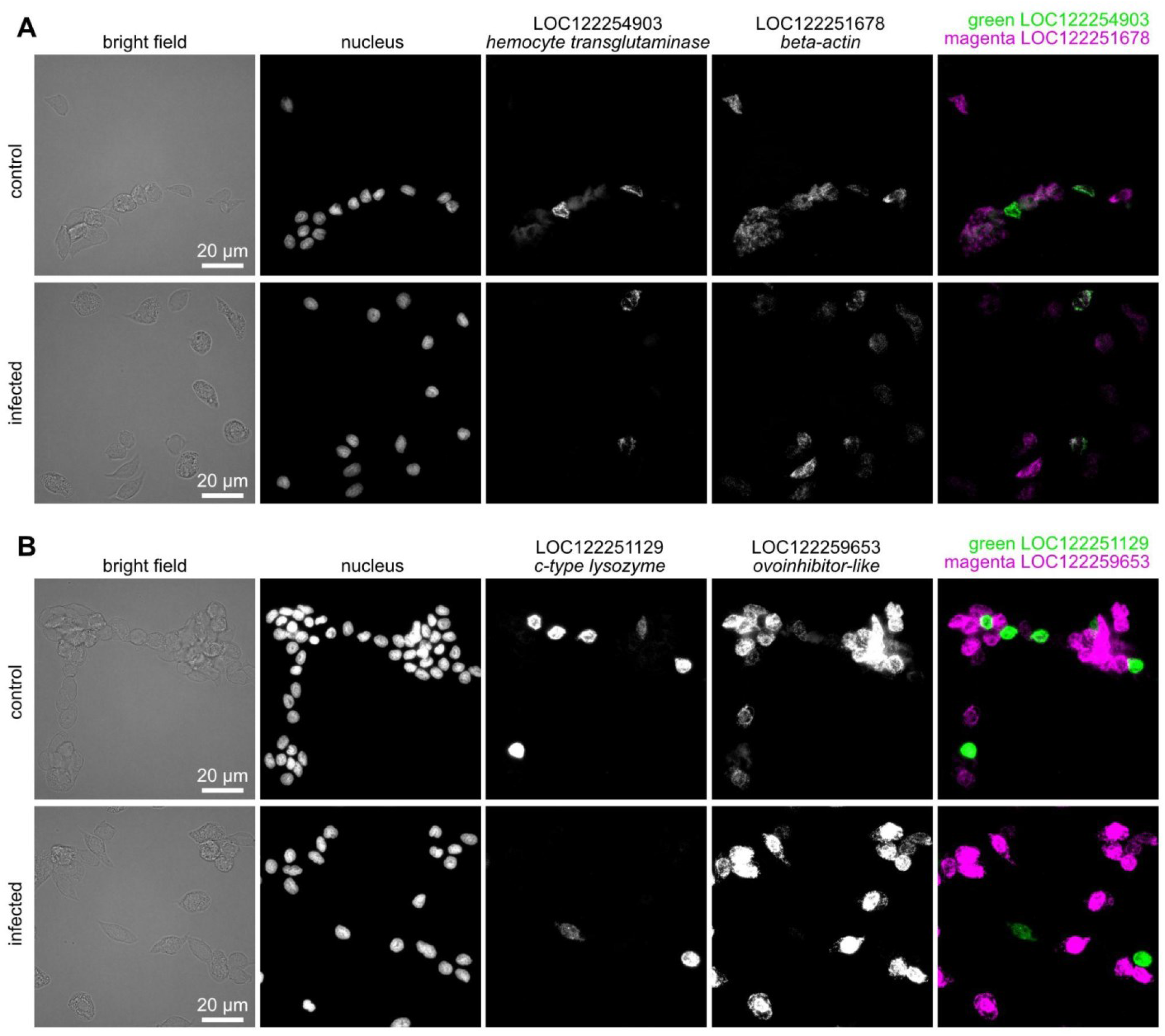
Detection of transcripts of candidate marker genes by single molecule fluorescence *in situ* hybridization (smFISH). Transcript detection results for the positive control LOC122251678 *beta-actin* and Hem1 marker candidate LOC122254903 *hemocyte transglutaminase* (A). Transcript detection results for Hem4 marker candidate LOC122259653 *ovoinhibitor-like*, and Hem7 marker candidate LOC122251129 *c-type lysozyme* (B)

### Estimation of new antimicrobial peptides from gene expression patterns

Antimicrobial peptides (AMPs) were highly expressed in cluster Hem4-5 and cluster Hem7 (Figure 6). LOC122251298 anti-lipopolysaccharide factor-like and LOC122260788 arasin 1-like were upregulated in cluster Hem7 by viral infection. On the other hand, AMP expression in the cluster Hem4-5 was decreased by viral infection. Therefore, we extracted a gene whose function was unknown, which was expressed in specific clusters like other AMPs, and whose expression was reduced due to viral infection, and evaluated the possibility that this gene is an AMP. The deduced amino acid sequences of these unknown genes were estimated and examined for possible AMPs by the Antimicrobial Peptide Scanner vr.2 (Veltri et al., 2018) and AI4AMP (Lin et al., 2021). As a result, LOC122242927 and LOC122250143 were predicted as AMPs. LOC122242927 could not be predicted by either domain search; the Antimicrobial Peptide Scanner vr.2 showed 0.9999 probability and the AI4AMP showed maximum 0.8262734 probability of AMP; ESMFold predicted a β-sheet structure at the N-terminal and an α-helix structure at the C-terminal (Figure 7A). LOC122250143 could not be predicted by any of the domain searches; the Antimicrobial Peptide Scanner vr.2 showed 1.0 probability and the AI4AMP showed maximum 0.9984921 probability of AMP; ESMFold predicted a linear extension structure without predicting any of the folding (Figure 7B). On their surfaces, both proteins were expected to have cationic charges and hydrophobic amino acids (Figure 7CD).

**Figure 6.**
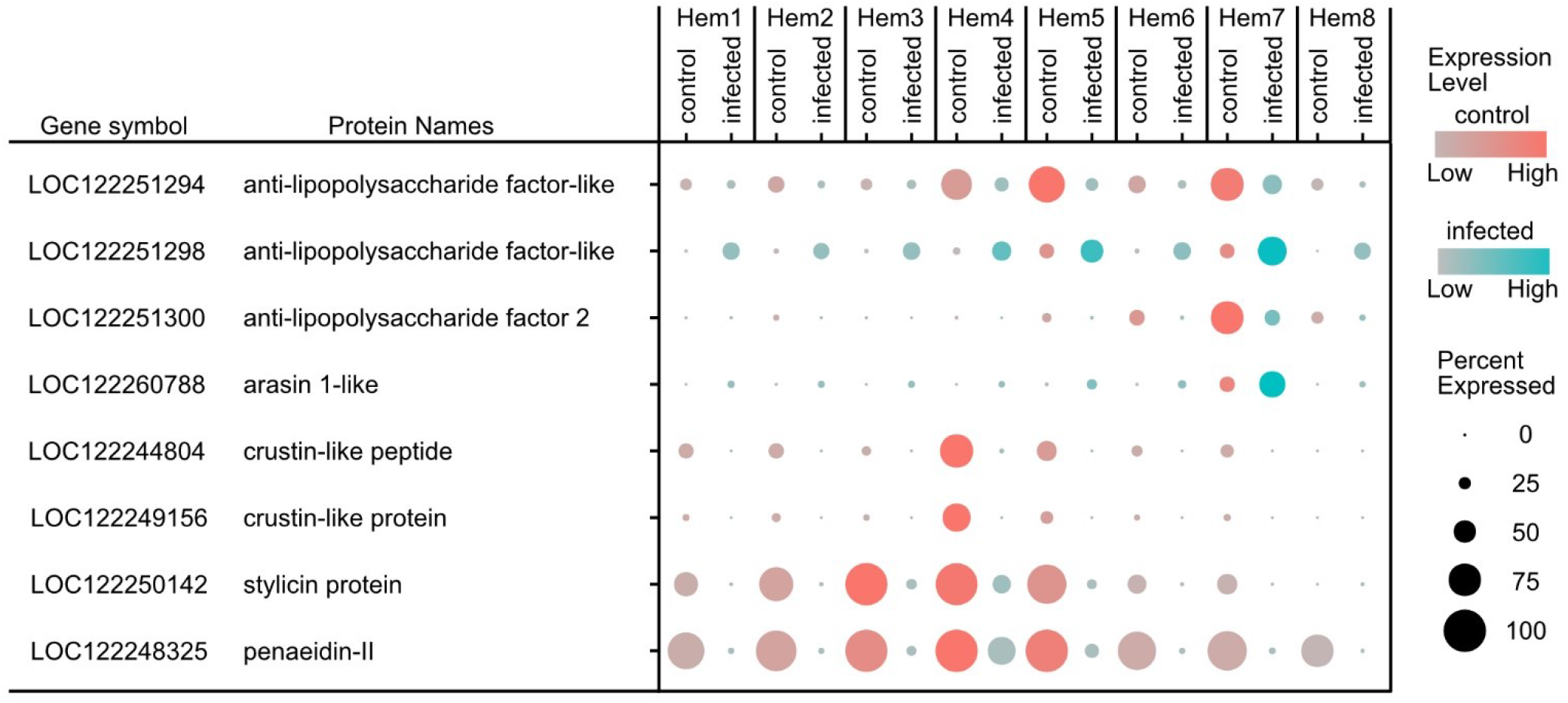
Expression patterns of antimicrobial peptides (AMPs) detected in this study in hemocytes of *P. japonicus*. The expression of AMPs was centered in clusters Hem4,5,7, and also found to be reduced by WSSV infections, except for LOC122251298 and LOC122260788.

**Figure 7.**
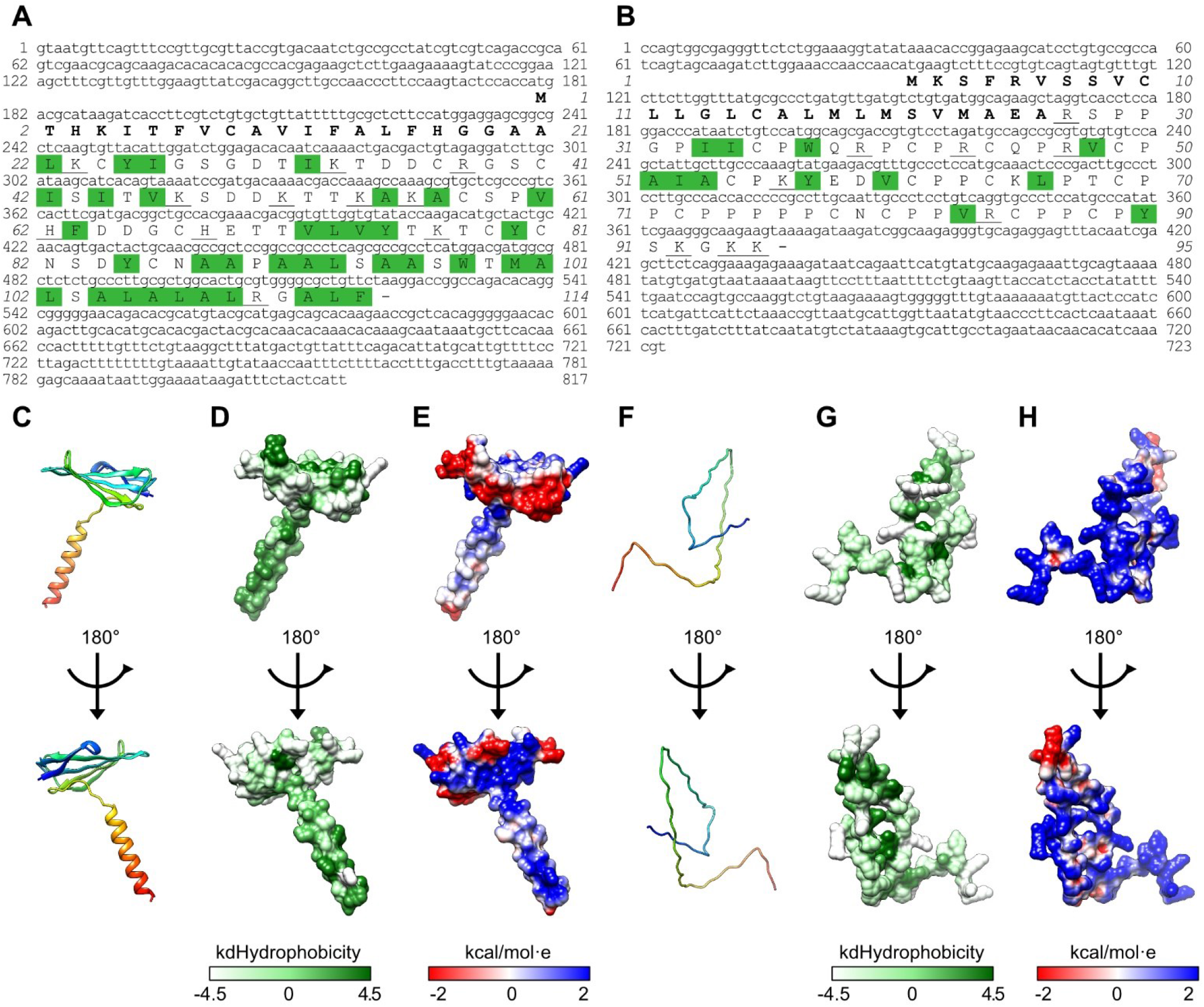
The sequences structures, and the visualizations of surface hydrophobicity and electrostatic properties of candidate antimicrobial peptides LOC122242927 and LOC12250143. The transcribed sequence and deduced amino acids sequence of LOC122242927 (A) and LOC122250143 (B). Bold amino acids represent estimated signal peptides, underlines under an amino acid indicates that the amino acid has a cationic charge, green highlight indicates that the amino acid is hydrophobic. The ribbon style amino acid structure of LOC122242927 (C) and LOC122250143 (F).The surface representation of hydrophobicity of LOC122242927 (D) and LOC122250143 (G) and electrostatic properties of LOC122242927 (E) and LOC122250143 (H). The hydrophobicity properties of amino acids are indicated using the Kyte and Doolittle hydrophobicity scale; the most polar residues are in white and the most hydrophobic residues are in dark green in the surface representation. The electrostatic potential ranges from negative (red) to positive (blue). The structures of both amino acids were displayed using the surface coloring feature of the UCSF Chimera tool (v.1.16).

## Discussion

### The effect of methanol fixing of hemocytes for scRNA-seq

The methanol fixation decreases the efficiency of the UMI and gene detection per cell. However, since UMAP analysis and marker gene detection can be performed, rough gene transcription in a cell population can be tracked. The advantage of methanol fixation to avoid performing scRNA-seq immediately after sampling is convenient for non-model organisms, where the sampling site and the experimental site are often different. On the other hand, Drop-seq is less sensitive to genes and UMI than HyDrop or 10X Chromium, which perform RT in the droplet, because RT is performed after droplet breakage. Of course, it would be desirable to be able to generate all data from HyDrop or 10X Chromium, but this is difficult due to the versatility of the equipment and the high cost of reagents. When conducting single-cell analysis on non-model organisms, first run scRNA-seq, which can analyze a lot of cells with high sensitivity, like HyDrop or 10X Chromium, to figure out how the entire cell population will be distributed. Then, increasing the sample size and monitoring changes in cell populations should be accomplished using affordable techniques like Drop-seq.

### Viral response populations and their gene expression changes

Previous studies have shown that WSSV infection decreases the number of circulating hemocytes by a factor of ten or more, from 10^7^ to 10^5^ hemocytes/mL in hemolymph (Cui et al., 2020; Elbahnaswy et al., 2017; van de Braak et al., 2002; Wongprasert et al., 2003), that causes a serious breakdown of the immune system, leading to the death of shrimp. WSSV infects specific types of hemocytes (Wang et al., 2002) and WSSV causes changes in the expression level of immune-related genes (Wang et al., 2019; Xue et al., 2013). In this study, genes such as LOC122260926 *syndecan-like*, which had an increase in gene expression by virus infection in total hemocytes, had also previously been reported to have similar increases (Sun et al., 2014; Yang et al., 2015). It is also interesting to note that the syndecan is thought to be involved in the entry of WSSV into cells, and that the expression of these receptors has increased due to viral infection. On the other hand, *antimicrobial peptides* (*AMPs*) such as LOC122248325 *penaeidin-II* and LOC122250142 *stylicin protein* had decreased in expression due to virus infection. It is known that AMPs are expressed in granulocytes, and they are discussed as markers for defining granulocytes. Zhang (Zhang et al., 2018) used antibodies against penaeidin-II to perform immune staining on hemocytes and found that the number of hemocytes expressing penaeidin-II strongly decreased in virus-infected individuals compared to healthy individuals at the protein level. However, due to the lack of markers at this point, it was only possible to discuss that the populations expressing AMPs had disappeared. This study has shown that the cluster Hem4-centered population does indeed decrease in proportion as a whole due to viral infection, but it still exists as a population and has become unable to express AMPs. Although the changing of the cell population in the circulating hemolymph was analyzed, it has been reported that hemocytes migrate to seriously infected tissue when infected with a virus (van de Braak et al., 2002; Wongprasert et al., 2003). Therefore, from the results of this study, it is suggested that clusters of hemocytes that express AMPs in the circulating hemolymph migrate to seriously infected tissue during infection with pathogens, resulting in a decrease in the apparent number of hemocytes in the circulating hemolymph. While analysis of tissue sections using cell markers other than AMPs is also necessary, continuing single-cell analysis using cells prepared from tissue will be necessary to study in depth.

In addition to AMPs, *single IGFBPs* were highly expressed in cluster Hem4, and their expression was decreased during viral infection. These results indicate that this protein group is important for viral infection and the expression of AMPs. These single IGFBPs were shown to be homologous with other crustacean’s crustacean hematopoietic factors (CHFs). Silencing of CHF or its candidate receptor, laminin, has been reported to decrease the number of hemocytes, especially hyaline cells (Charoensapsri et al., 2015; Lin et al., 2011; Söderhäll, 2013). Recent studies suggest that granulocytes differentiate from hyaline cells; it is possible that viral infection suppresses CHF expression, resulting in cell death at the hyaline cell stage, reducing the cluster Hem4 and AMPs expression, or preventing the hemocyte differentiation from hyaline cells into hemocytes expressing AMPs (Cui et al., 2022; Koiwai et al., 2021; Li et al., 2021; Liu et al., 2021; Li et al., 2022). The critical function of CHFs remains unclear, but effective anti-WSSD control strategies may depend on the protection of AMPs-expressing populations by controlling CHFs expression.

### Advantages of scRNA-seq for exploring new markers and using them in non-model organisms

Previous studies did not have markers to define cell populations, so it was not clear whether the decrease in the expression of AMPs was due to a decrease in the number of AMPs-expressing populations or a decrease in expression within cell populations. While morphological classification is certainly important, because it is simple and easy, it is also important to discover marker genes that are unlikely to change in expression under any circumstances when defining and studying the function of each cell population in the future. These genes should be used to define cell types based on their expression. In this study, we identified marker genes and found that three of them function as markers: LOC122254903 *hemocyte transglutaminase* (HemTGase), LOC122259653 *ovoinhibitor-like* (OIH), and LOC122251129 *c-type lysozyme* (LYS), through smFISH. It will be necessary to classify hemocyte populations using these genes, with notations like HemTGase^+^ hemocytes, OIH^+^ hemocytes, or LYZ^+^ hemocytes. It has also been studied that AMPs are important for immune function, and it is possible to objectively measure and clarify why the expression of these AMPs decreases and how to stop the decrease in expression, which will be useful in countermeasures against viral diseases. In previous systems, it was not possible to perform these cell population analyses without specific antibodies for marker genes, and even if it was possible, only a few markers could be followed. The powerful part of scRNA-seq is that it can follow the changes in the expression of more than hundreds of genes at once in a single cell. This allows analysis of how cell populations and gene expression changed under specific conditions even in non-model organisms without protein markers.

### Prediction of functions of functionally unknown genes from the population analysis

We have identified two candidate genes for antimicrobial peptides, LOC122242927 and LOC122250143. LOC122242927 has a β-sheet structure at the N-terminal and an α-helix structure at the C-terminal, respectively, while neither structure was predicted for LOC122250143. AMPs are known to be simultaneously having an α-helix and a β-sheet; others have a linear extension structure in aqueous solution and fold a specific conformation in bacterial membranes (Huan et al., 2020). Each of the newly identified AMP candidates is presumed to be one of these types.

In recent decades, genome sequencing has become more common, and transcriptome sequencing has been combined with it to predict genes using bioinformatics. However, in the standard settings of many gene prediction tools, sequences shorter than 100 amino acid residues are labeled as non-coding genes. As a result, functional groups of short proteins such as AMPs are missed. In addition, the function of unknown genes is often named based on the domains they contain in their sequence, but it is difficult to predict the function of genes without any domains. Therefore, in this study, we focused on the cell population expressing the gene rather than the gene itself. Paradoxically, we hypothesized that unknown genes expressed in specific cell populations with specific functions may have the same functions, and were able to identify novel AMP candidate genes. This approach to single-cell analysis is new and is expected to be widely used in non-model organisms with many unknown genes.

### Summary

In this study, we were able to analyze the dynamics of hemocyte populations without using protein markers by performing single-cell RNA analysis on multiple individuals. By fixing the target cells with methanol, we were able to multiplex the target samples and improve the efficiency of the workflow. It was found that cluster Hem4 decreased due to viral infection. To protect against viral diseases, it seems advisable to suppress the decrease in cluster Hem4. In addition to suppressing the decrease in AMPs, which directly act on pathogenic bacteria, research focusing on CHFs, which are thought to control the differentiation of cluster Hem4, will be important. Single-cell analysis can not only classify cell populations, but also infer the functions of unknown genes expressed in cell populations with certain functions. By continuing to analyze hemocytes subjected to stress or other pathogenic bacteria, the classification and functional analysis of this cell population will continue.

## Material and method

### Shrimp and single-cell suspension for Drop-seq

Twenty-to twenty-five-gram shrimp were purchased from a local farmer and maintained in artificial seawater with 30-35 ppt salinity with a recirculating system at 25 ^0^C. Hemolymph was collected using an anticoagulant solution suitable for penaeid shrimp from an abdominal site. The collected hemolymph was centrifuged at 800 x *g* for 10 min to collect the hemocytes, which were then washed twice with PBS and the osmolarity was adjusted to kuruma shrimp (KPBS: 480 mM NaCl, 2.7 mM KCl, 8.1 mM Na2HPO4·12H2O, 1.47 mM KH2PO4, pH 7.4). The hemocyte concentration was adjusted to 5 × 10^6^ cells/mL using 0.05% BSA in KPBS. A final methanol concentration of 80% was used to fix these adjusted hemocytes. Prior to use, the fixed hemocytes were kept in a freezer at -80 ^0^C.

### Artificial infection experiment

Since *in vitro* culture of the WSSV is not possible, WSSV-infected shrimp individuals were homogenized and filtered to use the solution as the source of the WSSV infection solution. The copy number of WSSV in the infection source was quantified by real-time PCR with direct lysis buffer (Li et al., 2011; Sun et al., 2013). Artificial infection was performed using this infection solution, and the copy number of WSSV in the gills was quantified at the same time as a sampling of hemocytes. Individuals with more than 1 × 10^5^ WSSV copies/ng of total genomic DNA were subjected to analysis by Drop-seq.

### Drop-seq

The Drop-seq procedure was used to encapsulate single hemocytes and single mRNA capture beads into fL-scale microdroplets, as previously described (Macosko et al., 2015). Briefly, the self-built Drop-seq microfluidic device was prepared by molding polydimethylsiloxane (PDMS; Sylgard 184, Dow Corning Corp.) from the microchannel structure using the negative photoresist (SU-8 3050, Nippon Kayaku Co.). Using this device, droplets containing a cell and a Barcoded Bead SeqB (ChemGenes Corporation) were produced up to 2 mL per sample using a pressure pump system (On-chip Droplet generator, On-chip Biotechnologies Co., Ltd.). During the sample introduction, the vial bottles containing cells and beads were shaken using a vortex mixer to prevent sedimentation and aggregation (Biočanin et al., 2019). The single suspension of hemocytes was prepared from methanol-fixed hemocytes as follows, the fixed hemocytes stored at -80 ^0^C were centrifuged at 1,000 × *g* for 5 min, then washed once with 3 × SSC buffer containing 0.04% BSA, 200 U/mL of RNase inhibitor and 1 mM DTT, the number of hemocytes was set as 1.2 × 10^5^ cells/buffer. Droplets were collected from the channel outlet into the 50 mL conical tube. Then, droplets were broken promptly and barcoded beads with captured transcriptomes were reverse transcribed using Maxima H Minus Reverse Transcriptase (Thermo Fisher Scientific) at 25 ^0^C for 30 min, then at 42 ^0^C for 90 min. Then, the beads were treated with Exonuclease I (New England Biolabs) to obtain single-cell transcriptomes attached to microparticles (STAMP). The first-strand cDNAs on beads were amplified using PCR. The beads obtained above were distributed throughout PCR tubes (2,000 beads per tube), wherein 1 × KAPA HiFi HS Ready Mix (KAPA Biosystems) and 0.8 μM 1st PCR primer were included in a 50 µL reaction volume. PCR amplification was achieved using the following program: initial denaturation at 95 ^0^C for 3 min; 4 cycles at 98 ^0^C for 20 s, 65 ^0^C for 45 s, and 72 ^0^C for 6 min; 11 cycles of 98 ^0^C for 20 s, 67 ^0^C for 20 s, and 72 ^0^C for 6 min; and a final extension at 72 ^0^C for 5 min. The amplicons were pooled, double-purified with × 0.6 SeraMag (Cytiva), and eluted in 35 µL of ddH2O. Sequence-ready libraries were prepared using the NexteraXT kit (Illumina). A total of 600 pg of each cDNA library was fragmented using transposome. The amplified library was purified using × 0.6 once continuously × 1.0 SeraMag beads and sequenced (paired-end) on an Illumina NovaSeq 600 sequencer (SP flowcell); 20 cycles for read1 with custom sequence primer, 8 cycles for index read, and 102 cycles for read2.

### Analysis of single-cell data

Paired-end reads were processed and mapped to the genome sequence of *P. japonicus* (Kawato et al., 2021) by STARsolo (Dobin et al., 2013). After the digital expression data of eight shrimps from this study and three from the previous study (Koiwai et al., 2021) were calculated, a total of 11 data were read by Seurat v4.1.1. The single-cell data were extracted with the following parameters: a number of UMIs per cell greater than 500; a number of UMIs per cell less than 4,000; and a percentage of mitochondrial genes less than 10%. Then, SCTransform V2 was performed to remove the technical variation while retaining biological heterogeneity. We ran a PCA using the expression matrix of the top 3,000 most variable genes. The total number of principal components (PCs) required to compute and store was 30. The UMAP was then performed using the following parameters: n.neighbors, min.dist and n.components were set to 30, 0.3, and 2, respectively, to visualize the data in the two-dimensional space, and then the clusters were predicted with a resolution of 0.5. Up- and down-regulated genes by WSSV infection of each cluster were predicted by FindMarkers function of Seurat with the parameters: min.pct and logfc.threshold were 0.5 and 1.0 using SCT assay slot. Conserved markers were predicted by the program FindConservedMarkers of Seurat, with the parameters: min.pct and logfc.threshold were 0.5 and 1.0, respectively.

### mRNA detection of conserved markers by smFISH

The genes predicted as conserved markers were validated by smFISH. The smFISH probes against each gene were designed from the genome sequence of *P. japonicus*. The hemolymph was withdrawn using an anticoagulant, then washed twice with KPBS. Hemocytes were fixed by 10% formalin in KPBS for 15 min at RT. The fixed hemocytes suspension was spread on a cover glass (No.1) round Φ18mm in a cell collection bucket SC-2 (TOMY) at 100 × *g* for 2 min, then permeabilized by 80% EtOH for 60 min at 4 ^0^C. The probes were hybridized overnight at 37 ^0^C in the hybridization buffer (10% w/v dextran sulfate and 10% formamide in 2 × SSC). After hybridization, cells were washed with wash buffer (10% formamide in 2 × SSC) twice, then stained the nucleus by 1 μg/mL of Hoechst 33342 (DOJINDO LABORATORIES). The stained hemocytes were subjected to an inverted microscope (BZ-X810: KEYENCE CORPORATION) using suitable filter sets. To obtain the maximum smFISH signal, images were taken at a pitch of 0.1 μm on the y-axis and combined. Then the positive number of hemocytes was counted using Fiji (Schindelin et al., 2012). Custom Stellaris® FISH Probes were designed against LOC122251678 *beta-actin*, LOC122254903 *hemocyte transglutaminase*, LOC122259653 *ovoinhibitor-like*, and LOC122251129 *c-type lysozyme* by utilizing the Stellaris® RNA FISH Probe Designer (Biosearch Technologies, Inc., Petaluma, CA) available online at www.biosearchtech.com/stellarisdesigner (Version 4.2). Stellaris RNA FISH Probe set labeled with Quasar 570 or 670 dye (Biosearch Technologies, Inc.), following the manufacturer’s instructions available online at www.biosearchtech.com/stellarisprotocols. The number of analyzed hemocytes was 115 (control) and 151 (infected) for detecting LOC122254903 and LOC122251678, and 162 (control) and 144 (infected) for detecting LOC12225963 and LOC122251129, respectively.

### Estimation of new antimicrobial peptides from functionally unknown genes

Antimicrobial peptides of kuruma shrimp were extracted from its genome sequence following a review paper (Saucedo-Vázquez et al., 2022; Tassanakajon et al., 2013; Tincu & Taylor, 2004). Eight of thirty-four genes were analyzed on this Drop-seq. Functionally unknown genes that showed the same expression pattern in each cluster with and without virus infection were manually extracted. The existence of a signal peptide sequence was predicted by signaIP version 6.0 (Teufel et al., 2022). The possibility of AMP in functionally unknown genes without a signal peptide region was predicted by the Antimicrobial Peptide Scanner vr.2 (Veltri et al., 2018) and AI4AMP (Lin et al., 2021). The structures of the deduced amino acids sequences were predicted by ESMFold, then the structures were visualized using UCSF Chimera 1.16.

### Statistics

The Wilcoxon signed-rank test and one-way analysis of variance were performed on R software.

### Data files and analysis code

The raw sequence data of newly sequenced *P. japonicus* Drop-seq transcriptomic reads were archived in the DDBJ Sequence Read Archive (DRA) as DRA015407 under the DNA Data Bank of Japan.

## Acknowledges

We would like to thank On-chip Biotechnologies Co., Ltd. for providing their pressure pump On-chip Droplet Generator, Hiroaki Suzuki (Department of Precision Mechanics, Faculty of Science and Engineering, Chuo University, Bunkyo, Japan) and Ryuji Kawano (Department of Biotechnology and Life Science, Tokyo University of Agriculture and Technology, Koganei, Japan) for their technical support in the fabrication of the Drop-seq microfluidic devices, and GenomeLead for the sequencing of Drop-seq libraries by NovaSeq 600. This work was supported by Japan Society for Promotion of Science (JSPS) KAKENHI Grant-in-Aid for Early-Career Scientists Grant Number 20K15603 and Japan Science and Technology Agency (JST) ACT-X Grant Number JPMJAX21B5 to Keiichiro Koiwai; JSPS KAKENHI Grant Numbers 22H00379 and Science and Technology Research Partnership for Sustainable Development (SATREPS) in collaboration between JST and Japan International Cooperation Agency (JICA) Grant Number JPMJSA1806 to Ikuo Hirono.

## Author contributions

K. Koiwai designed the experiments, performed the experiments, analyzed the data, and wrote the manuscript; H. Kondo and I. Hirono supervised the research.

## Reference

Alles, J., Karaiskos, N., Praktiknjo, S. D., Grosswendt, S., Wahle, P., Ruffault, P. L., Ayoub, S., Schreyer, L., Boltengagen, A., Birchmeier, C., Zinzen, R., Kocks, C., & Rajewsky, N. (2017). Cell fixation and preservation for droplet-based single-cell transcriptomics. BMC Biology, 15(1), 44. https://doi.org/10.1186/s12915-017-0383-5

Biočanin, M., Bues, J., Dainese, R., Amstad, E., & Deplancke, B. (2019). Simplified Drop-seq workflow with minimized bead loss using a bead capture and processing microfluidic chip. Lab on a Chip, 19(9), 1610–1620. https://doi.org/10.1039/c9lc00014c

Charoensapsri, W., Sangsuriya, P., Lertwimol, T., Gangnonngiw, W., Phiwsaiya, K., & Senapin, S. (2015). Laminin receptor protein is implicated in hemocyte homeostasis for the whiteleg shrimp Penaeus (Litopenaeus) vannamei. Developmental and Comparative Immunology, 51(1), 39–47. https://doi.org/10.1016/j.dci.2015.02.012

Cui, C., Liang, Q., Tang, X., Xing, J., Sheng, X., & Zhan, W. (2020). Differential apoptotic responses of hemocyte subpopulations to white spot syndrome virus infection in Fenneropenaeus chinensis. Frontiers in Immunology, 11, 594390. https://doi.org/10.3389/fimmu.2020.594390

Cui, C., Tang, X., Xing, J., Sheng, X., Chi, H., & Zhan, W. (2022). Single-cell RNA-seq uncovered hemocyte functional subtypes and their differentiational characteristics and connectivity with morphological subpopulations in Litopenaeus vannamei. Frontiers in Immunology, 13, 980021. https://doi.org/10.3389/fimmu.2022.980021

De Rop, F., Ismail, J. N., Bravo González-Blas, C., Hulselmans, G. J., Flerin, C. C., Janssens, J., Theunis, K., Christiaens, V. M., Wouters, J., Marcassa, G., de Wit, J., Poovathingal, S., & Aerts, S. (2022). HyDrop enables droplet based single-cell ATAC-seq and single-cell RNA-seq using dissolvable hydrogel beads. eLife, 11. https://doi.org/10.7554/eLife.73971

Dobin, A., Davis, C. A., Schlesinger, F., Drenkow, J., Zaleski, C., Jha, S., Batut, P., Chaisson, M., & Gingeras, T. R. (2013). STAR: ultrafast universal RNA-seq aligner. Bioinformatics, 29(1), 15–21. https://doi.org/10.1093/bioinformatics/bts635

Elbahnaswy, S., Koiwai, K., Zaki, V. H., Shaheen, A. A., Kondo, H., & Hirono, I. (2017). A novel viral responsive protein (MjVRP) from Marsupenaeus japonicus haemocytes is involved in white spot syndrome virus infection. Fish & Shellfish Immunology, 70, 638–647. https://doi.org/10.1016/j.fsi.2017.09.045

Flegel, T. W. (2012). Historic emergence, impact and current status of shrimp pathogens in Asia. Journal of Invertebrate Pathology, 110(2), 166–173. https://doi.org/10.1016/j.jip.2012.03.004

Flegel, T. W. (2019). A future vision for disease control in shrimp aquaculture. Journal of the World Aquaculture Society, 50(2), 249–266. https://doi.org/10.1111/jwas.12589

Huan, Y., Kong, Q., Mou, H., & Yi, H. (2020). Antimicrobial peptides: classification, design, application and research progress in multiple fields. Frontiers in Microbiology, 11, 582779. https://doi.org/10.3389/fmicb.2020.582779

Kawato, S., Nishitsuji, K., Arimoto, A., Hisata, K., Kawamitsu, M., Nozaki, R., Kondo, H., Shinzato, C., Ohira, T., Satoh, N., Shoguchi, E., & Hirono, I. (2021). Genome and transcriptome assemblies of the kuruma shrimp, Marsupenaeus japonicus. G3, 11(11). https://doi.org/10.1093/g3journal/jkab268

Klein, A. M., Mazutis, L., Akartuna, I., Tallapragada, N., Veres, A., Li, V., Peshkin, L., Weitz, D. A., & Kirschner, M. W. (2015). Droplet barcoding for single-cell transcriptomics applied to embryonic stem cells. Cell, 161(5), 1187–1201. https://doi.org/10.1016/j.cell.2015.04.044

Koiwai, K., Koyama, T., Tsuda, S., Toyoda, A., Kikuchi, K., Suzuki, H., & Kawano, R. (2021). Single-cell RNA-seq analysis reveals penaeid shrimp hemocyte subpopulations and cell differentiation process. eLife, 10, e66954. https://doi.org/10.7554/eLife.66954

Li, F., Zheng, Z., Li, H., Fu, R., Xu, L., & Yang, F. (2021). Crayfish hemocytes develop along the granular cell lineage. Scientific Reports, 11(1), 13099. https://doi.org/10.1038/s41598-021-92473-9

Li, H., Xu, H., Zhao, C., Sulaiman, Y., & Wu, C. (2011). A PCR amplification method without DNA extraction. Electrophoresis, 32(3-4), 394–397. https://doi.org/10.1002/elps.201000392

Lin, T.-T., Yang, L.-Y., Lu, I.-H., Cheng, W.-C., Hsu, Z.-R., Chen, S.-H., & Lin, C.-Y. (2021). AI4AMP: an Antimicrobial peptide predictor using physicochemical property-based encoding method and deep learning. mSystems, 6(6), e0029921. https://doi.org/10.1128/mSystems.00299-21

Lin, X., Söderhäll, K., & Söderhäll, I. (2011). Invertebrate hematopoiesis: an astakine-dependent novel hematopoietic factor. Journal of Immunology, 186(4), 2073–2079. https://doi.org/10.4049/jimmunol.1001229

Lin, Z., Akin, H., Rao, R., Hie, B., Zhu, Z., Lu, W., Smetanin, N., Verkuil, R., Kabeli, O., Shmueli, Y., dos Santos Costa, A., Fazel-Zarandi, M., Sercu, T., Candido, S., & Rives, A. (2022). Evolutionary-scale prediction of atomic level protein structure with a language model. In bioRxiv (p. 2022.07.20.500902). https://doi.org/10.1101/2022.07.20.500902

Liu, M.-J., Liu, S., & Liu, H.-P. (2021). Recent insights into hematopoiesis in crustaceans. Fish and Shellfish Immunology Reports, 2, 100040. https://doi.org/10.1016/j.fsirep.2021.100040

Li, Y., Zhou, F., Yang, Q., Jiang, S., Huang, J., Yang, L., Ma, Z., & Jiang, S. (2022). Single-cell sequencing reveals types of hepatopancreatic cells and haemocytes in black tiger shrimp (Penaeus monodon) and their molecular responses to ammonia stress. Frontiers in Immunology, 13. https://doi.org/10.3389/fimmu.2022.883043

Macosko, E. Z., Basu, A., Satija, R., Nemesh, J., Shekhar, K., Goldman, M., Tirosh, I., Bialas, A. R., Kamitaki, N., Martersteck, E. M., Trombetta, J. J., Weitz, D. A., Sanes, J. R., Shalek, A. K., Regev, A., & McCarroll, S. A. (2015). Highly parallel genome-wide expression profiling of individual cells using nanoliter droplets. Cell, 161(5), 1202–1214. https://doi.org/10.1016/j.cell.2015.05.002

Meng, J., & Wang, W.-X. (2022). Highly sensitive and specific responses of oyster hemocytes to copper exposure: single-cell transcriptomic analysis of different cell populations. Environmental Science & Technology, 56(4), 2497–2510. https://doi.org/10.1021/acs.est.1c07510

Meng, J., Zhang, G., & Wang, W.-X. (2022). Functional heterogeneity of immune defenses in molluscan oysters Crassostrea hongkongensis revealed by high-throughput single-cell transcriptome. Fish & Shellfish Immunology, 120, 202–213. https://doi.org/10.1016/j.fsi.2021.11.027

Naylor, R. L., Hardy, R. W., Buschmann, A. H., Bush, S. R., Cao, L., Klinger, D. H., Little, D. C., Lubchenco, J., Shumway, S. E., & Troell, M. (2021). A 20-year retrospective review of global aquaculture. Nature, 591(7851), 551–563. https://doi.org/10.1038/s41586-021-03308-6

Saucedo-Vázquez, J. P., Gushque, F., Vispo, N. S., Rodriguez, J., Gudiño-Gomezjurado, M. E., Albericio, F., Tellkamp, M. P., & Alexis, F. (2022). Marine arthropods as a source of antimicrobial peptides. Marine Drugs, 20(8). https://doi.org/10.3390/md20080501

Schindelin, J., Arganda-Carreras, I., Frise, E., Kaynig, V., Longair, M., Pietzsch, T., Preibisch, S., Rueden, C., Saalfeld, S., Schmid, B., Tinevez, J.-Y., White, D. J., Hartenstein, V., Eliceiri, K., Tomancak, P., & Cardona, A. (2012). Fiji: an open-source platform for biological-image analysis. Nature Methods, 9(7), 676–682. https://doi.org/10.1038/nmeth.2019

Söderhäll, I. (2013). Recent advances in crayfish hematopoietic stem cell culture: a model for studies of hemocyte differentiation and immunity. Cytotechnology, 65(5), 691–695. https://doi.org/10.1007/s10616-013-9578-y

Söderhäll, I., Fasterius, E., Ekblom, C., & Söderhäll, K. (2022). Characterization of hemocytes and hematopoietic cells of a freshwater crayfish based on single-cell transcriptome analysis. iScience, 104850. https://doi.org/10.1016/j.isci.2022.104850

Sun, X., Li, L., Wu, B., Ge, J., Zheng, Y., Yu, T., Zhou, L., Zhang, T., Yang, A., & Liu, Z. (2021). Cell type diversity in scallop adductor muscles revealed by single-cell RNA-Seq. Genomics. https://doi.org/10.1016/j.ygeno.2021.08.015

Sun, Y., Li, F., & Xiang, J. (2013). Analysis on the dynamic changes of the amount of WSSV in Chinese shrimp Fenneropenaeus chinensis during infection. Aquaculture, 376-379, 124–132. https://doi.org/10.1016/j.aquaculture.2012.11.014

Sun, Z., Li, S., Li, F., & Xiang, J. (2014). Bioinformatic prediction of WSSV-host protein-protein interaction. BioMed Research International, 2014, 416543. https://doi.org/10.1155/2014/416543

Tassanakajon, A., Somboonwiwat, K., Supungul, P., & Tang, S. (2013). Discovery of immune molecules and their crucial functions in shrimp immunity. Fish & Shellfish Immunology, 34(4), 954–967. https://doi.org/10.1016/j.fsi.2012.09.021

Teufel, F., Almagro Armenteros, J. J., Johansen, A. R., Gíslason, M. H., Pihl, S. I., Tsirigos, K. D., Winther, O., Brunak, S., von Heijne, G., & Nielsen, H. (2022). SignalP 6.0 predicts all five types of signal peptides using protein language models. Nature Biotechnology, 40(7), 1023–1025. https://doi.org/10.1038/s41587-021-01156-3

Tincu, J. A., & Taylor, S. W. (2004). Antimicrobial peptides from marine invertebrates. Antimicrobial Agents and Chemotherapy, 48(10), 3645–3654. https://doi.org/10.1128/AAC.48.10.3645-3654.2004

van de Braak, C. B. T., Botterblom, M. H. A., Huisman, E. A., Rombout, J. H. W. M., & van der Knaap, W. P. W. (2002). Preliminary study on haemocyte response to white spot syndrome virus infection in black tiger shrimp Penaeus monodon. Diseases of Aquatic Organisms, 51(2), 149–155. https://doi.org/10.3354/dao051149

Veltri, D., Kamath, U., & Shehu, A. (2018). Deep learning improves antimicrobial peptide recognition. Bioinformatics, 34(16), 2740–2747. https://doi.org/10.1093/bioinformatics/bty179

Wang, F., Li, S., Xiang, J., & Li, F. (2019). Transcriptome analysis reveals the activation of neuroendocrine-immune system in shrimp hemocytes at the early stage of WSSV infection. BMC Genomics, 20(1), 247. https://doi.org/10.1186/s12864-019-5614-4

Wang, X., Yu, L., & Wu, A. R. (2021). The effect of methanol fixation on single-cell RNA sequencing data. BMC Genomics, 22(1), 420. https://doi.org/10.1186/s12864-021-07744-6

Wang, Y. T., Liu, W., Seah, J. N., Lam, C. S., Xiang, J. H., Korzh, V., & Kwang, J. (2002). White spot syndrome virus (WSSV) infects specific hemocytes of the shrimp Penaeus merguiensis. Diseases of Aquatic Organisms, 52(3), 249–259. https://doi.org/10.3354/dao052249

Wongprasert, K., Khanobdee, K., Glunukarn, S. S., Meeratana, P., & Withyachumnarnkul, B. (2003). Time-course and levels of apoptosis in various tissues of black tiger shrimp Penaeus monodon infected with white-spot syndrome virus. Diseases of Aquatic Organisms, 55(1), 3–10. https://doi.org/10.3354/dao055003

Xue, S., Liu, Y., Zhang, Y., Sun, Y., Geng, X., & Sun, J. (2013). Sequencing and de novo analysis of the hemocytes transcriptome in Litopenaeus vannamei response to white spot syndrome virus infection. PloS One, 8(10), e76718. https://doi.org/10.1371/journal.pone.0076718

Yang, H., Li, S., Li, F., Wen, R., & Xiang, J. (2015). Analysis on the expression and function of syndecan in the Pacific white shrimp Litopenaeus vannamei. Developmental and Comparative Immunology, 51(2), 278–286. https://doi.org/10.1016/j.dci.2015.03.013

Yang, P., Chen, Y., Huang, Z., Xia, H., Cheng, L., Wu, H., Zhang, Y., & Wang, F. (2022). Single-cell RNA sequencing analysis of shrimp immune cells identifies macrophage-like phagocytes. eLife, 11. https://doi.org/10.7554/eLife.80127

Zhang, K., Koiwai, K., Kondo, H., & Hirono, I. (2018). White spot syndrome virus (WSSV) suppresses penaeidin expression in Marsupenaeus japonicus hemocytes. Fish & Shellfish Immunology, 78, 233–237. https://doi.org/10.1016/j.fsi.2018.04.045

Zhu, W., Yang, C., Chen, X., Liu, Q., Li, Q., Peng, M., Wang, H., Chen, X., Yang, Q., Liao, Z., Li, M., Pan, C., Feng, P., Zeng, D., & Zhao, Y. (2021). Single-cell ribonucleic acid sequencing clarifies cold tolerance mechanisms in the Pacific white shrimp (Litopenaeus vannamei). Frontiers in Genetics, 12, 792172. https://doi.org/10.3389/fgene.2021.792172

